# A unified framework for dopamine signals across timescales

**DOI:** 10.1101/803437

**Authors:** HyungGoo R. Kim, Athar N. Malik, John G. Mikhael, Pol Bech, Iku Tsutsui-Kimura, Fangmiao Sun, Yajun Zhang, Yulong Li, Mitsuko Watabe-Uchida, Samuel J. Gershman, Naoshige Uchida

## Abstract

Rapid phasic activity of midbrain dopamine neurons are thought to signal reward prediction errors (RPEs), resembling temporal difference errors used in machine learning. Recent studies describing slowly increasing dopamine signals have instead proposed that they represent state values and arise independently from somatic spiking activity. Here, we developed novel experimental paradigms using virtual reality that disambiguate RPEs from values. We examined the dopamine circuit activity at various stages including somatic spiking, axonal calcium signals, and striatal dopamine concentrations. Our results demonstrate that ramping dopamine signals are consistent with RPEs rather than value, and this ramping is observed at all the stages examined. We further show that ramping dopamine signals can be driven by a dynamic stimulus that indicates a gradual approach to a reward. We provide a unified computational understanding of rapid phasic and slowly ramping dopamine signals: dopamine neurons perform a derivative-like computation over values on a moment-by-moment basis.

## INTRODUCTION

Dopamine plays important roles in controlling learning, motivation, and movement. Understanding what types of information dopamine conveys is critical for determining how dopamine regulates various functions. One influential idea is that the activity of midbrain dopamine neurons represents temporal difference reward prediction errors (TD RPEs) used in reinforcement learning algorithms (Schultz et al., 1997; Daw et al., 2006; Niv, 2009; Ludvig et al., 2012; Gershman et al., 2014; Eshel et al., 2015; Starkweather et al., 2017; Watabe-Uchida et al., 2017). Dopamine neurons respond to an unpredicted reward or a reward-predicting cue with a short burst of spikes (or a ‘phasic’ excitation). Furthermore, omission of an expected reward causes a phasic inhibition of their firing. Besides these “phasic” responses, dopamine neurons typically maintain relatively low and stable baseline firing rates between these events. These response patterns have been observed in a number of animal species and in different task conditions (Bayer and Glimcher, 2005; Clark et al., 2012; Watabe-Uchida et al., 2017; Wenzel et al., 2015), and the RPE hypothesis has greatly impacted our understanding of dopamine functions in the brain. However, many of these experiments have employed relatively simple behavioral paradigms using discrete stimuli and outcomes. Whether the same principle applies to more complex situations remains to be examined.

Several studies that used freely moving animals have shown that dopamine concentrations in the striatum ramp up over the timescale of seconds (Phillips et al., 2003; Roitman et al., 2004; Howe et al., 2013; Hamid et al., 2016; Berke, 2018; Mohebi et al., 2019; Engelhard et al., 2019). Some authors have argued that these slow dopamine fluctuations cannot be readily explained by TD RPEs, and it has alternatively been proposed that they represent the value of the state (i.e., state value, or motivational value or value of work) that increases as the animal approaches a reward location (Berke, 2018; Hamid et al., 2016; Howe et al., 2013; Mohebi et al., 2019). Furthermore, a recent study (Mohebi et al., 2019) concluded that these ramping activities are absent in the spiking activity of dopamine neurons in the ventral tegmental area (VTA) and postulated that ramping dopamine signals arise from local modulations at their axon terminals in the striatum. Theoretically, it is thought that expected values, rather than RPEs, are well positioned to drive motivation (Berke, 2018). It is, thus, proposed that slowly fluctuating dopamine signals may underlie the role of dopamine in motivation (Berke, 2018; Hamid et al., 2016; Howe et al., 2013; Mohebi et al., 2019). However, more work is needed to determine (1) what mechanisms underlie the generation of ramping dopamine signals and (2) what behavioral conditions cause ramping dopamine signals.

Theoretically, value is unambiguously separable from RPE. TD RPE (δ*_t_*) is defined by

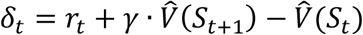

where *r_t_* is the reward that the animal receives at time *t*, *S_t_* is the state that the animal occupies at time 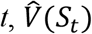 is the predicted value of the state *S_t_,* and *γ* is the discounting factor (*0 < γ < 1*) (Niv, 2009; Schultz et al., 1997). As evident from this equation, TD RPE contains the terms that are approximately the difference between the values of consecutive time points, *t* and 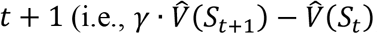 where *γ* is typically close to 1). In other words, TD RPEs are approximately the *derivative* of the value function as proposed originally (Sutton, 1988; Sutton and Barto, 1990) (see **Supplementary Note**). The idea that dopamine represents value is, therefore, incompatible with the canonical view that dopamine represents TD RPEs — the central idea in our current understanding of dopamine functions in reward processing.

Under many conditions, however, it has been difficult to disambiguate whether dopamine represents RPE or value. First, a dopamine ramp can occur regardless of whether dopamine represents value or RPEs (Gershman, 2014; Lloyd and Dayan, 2015; Morita and Kato, 2014). For instance, a theoretical study showed that the *shape* of the value function matters: if the value function is a *convex* function of proximity to reward, a TD RPE can exhibit a positive ramp (Gershman, 2014) (more general conditions are described in the **Supplementary Note**; see also Fig. S1). Therefore, the mere presence of a dopamine ramp does not distinguish the two possibilities. Second, value is a latent variable which must be inferred indirectly. A common approach has been to fit a model that contains value as a parameter and infer it from behavior (e.g. Hamid et al., 2016). These models, however, typically require various assumptions, leaving open the possibility that alternative models may better fit the data. Taken together, it remains unclear whether slowly fluctuating dopamine signals represent state values (motivational values) in the first place.

To circumvent these problems, here we sought to develop novel experimental paradigms that empirically dissociate RPE from value. Toward this goal, we focused on the core property of RPE, that is, RPE is approximately the derivative of value (**Supplementary Note**) (Doya, 2000; Schultz et al., 1997; Sutton, 1988; Sutton and Barto, 1990). Our experiments using visual virtual reality in mice allowed us to tease apart these two possibilities that are otherwise difficult to dissociate. The results show that ramping dopamine signals in the ventral striatum (nucleus accumbens) are consistent with TD RPEs but inconsistent with values. We also show that the spiking activity of VTA dopamine neurons as well as somatic calcium signals exhibit ramping activities that can explain ramping dopamine signals in the ventral striatum. Finally, we demonstrate that a non-navigational, simple visual cue that indicates a gradual increase in proximity to reward can also generate ramping dopamine signals. Together, our study demonstrates that the RPE account of dopamine signals can be extended to slowly fluctuating dopamine signals observed in dynamic sensory environments.

## RESULTS

### Experimental design: Using virtual reality to dissociate reward prediction errors from values

Before receiving reward, TD RPE is approximately the derivative of value, as discussed above. Basing on this core property of TD RPE, we sought to develop novel experimental paradigms that allow us to experimentally disambiguate value from RPE. Imagine that a mouse moves along a linear track to obtain reward (Fig. 1A). It is reasonable to assume that the value of location monotonically increases as the distance to the goal decreases. Now imagine that while moving in this linear track, the animal is suddenly *teleported* to a location closer to the goal (Fig. 1B). The value and RPE hypotheses make distinct predictions about the effect of this manipulation on the dopamine signal. If dopamine represents value, it should exhibit a step-like increase at the time of teleport and then continue increasing gradually, with the maximum level reached at the goal location (Fig. 1C, left). In stark contrast, if dopamine represents RPE, it should exhibit a phasic excitation at the time of teleport, reflecting an instantaneous increase in value (Fig. 1C, right). Because RPE depends on the *change* in value rather than the position per se, the level of dopamine need not be maximal at the goal and may also depend on the speed of movement across space.

**Fig. 1.**
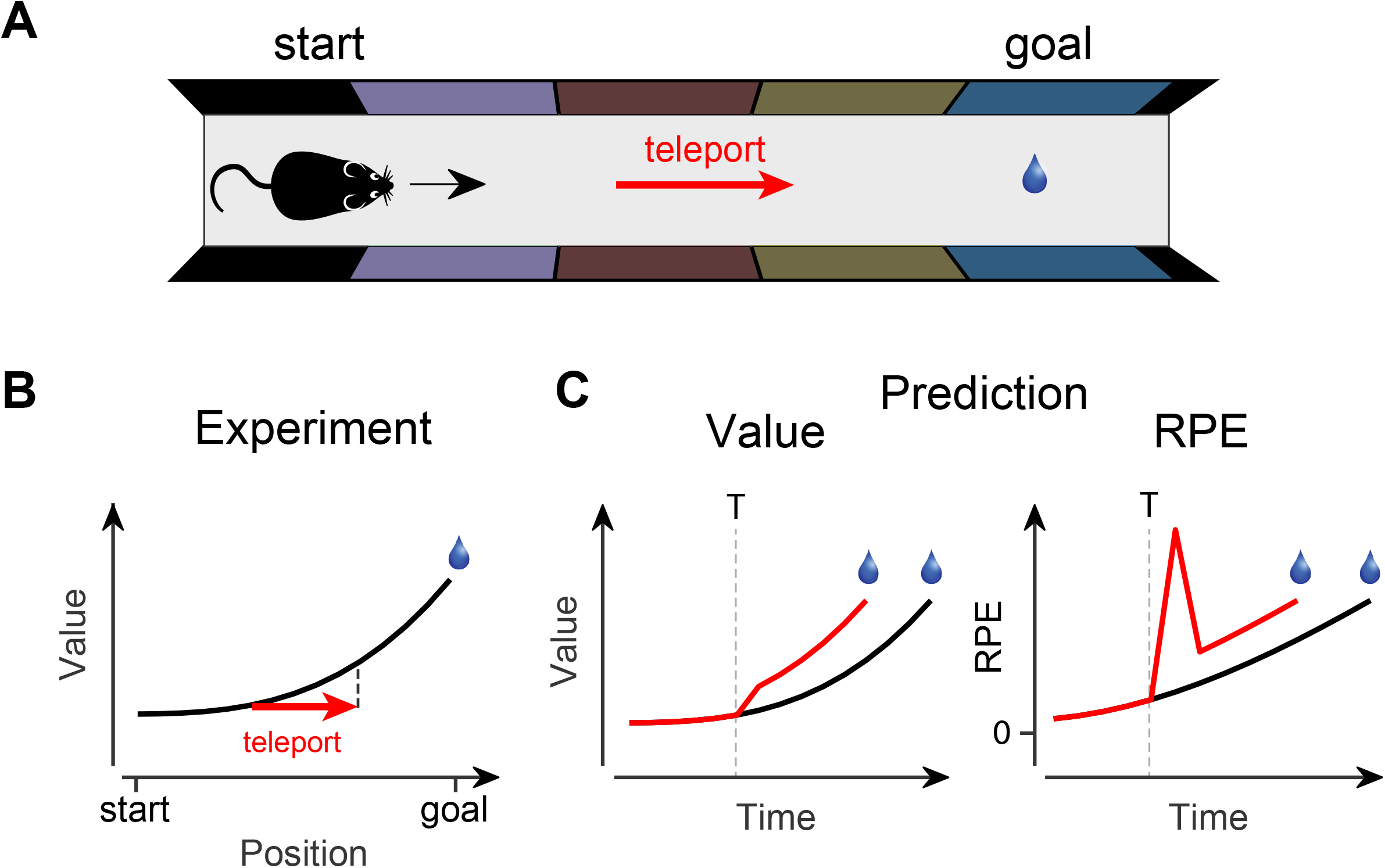
Experimental strategy to dissociate value versus reward prediction error (RPE) signals using teleport in visual virtual reality. **(A)** A top-down illustration of the virtual linear corridor. Different wall patterns indicated the current location to the animal. The animal was teleported from one location to another (red arrow). **(B)** The black curve indicates how the state value changes along the linear corridor. Red arrow: teleport. **(C)** Predictions of how the state value (left) and the TD RPE (right) are modulated by teleport (red curves). The value will exhibit a step-like increase, whereas TD RPE will exhibit a phasic excitation at the time of teleport.

In the present study, we empirically tested these ideas using visual virtual reality in head-fixed mice (Dombeck et al., 2007; Harvey et al., 2009) locomoting on a rotating wheel, which allowed us to manipulate the location of the animal, as determined by a visual scene, independently from its locomotion. The visual scene, which contained distinct wall patterns along the track (Fig. 2A), was moved at a constant speed, and the mice received a drop of water (5 µl) at the goal location (**Supplementary Movie 1**). Over several days of training, mice developed anticipatory licking near the goal location (Fig. 2B top, *n* = 16 mice, *P* = 0.00043, signed rank test). Although the visual scene was moved constantly irrespective of the animals’ behavior and locomotor movement was therefore not required for reward, more than half of the animals developed running behavior during the scene movement (Fig. 2B bottom).

**Fig. 2.**
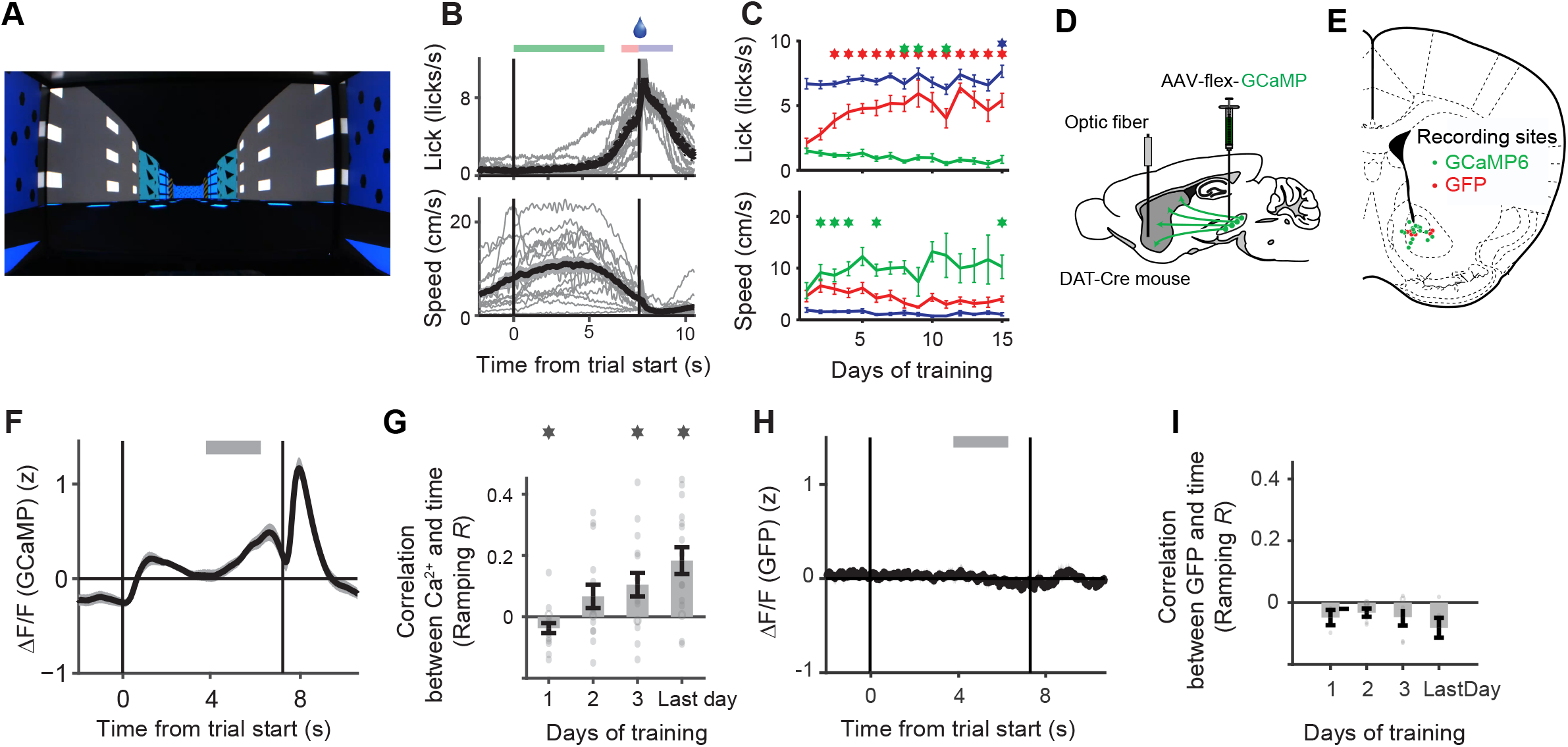
Behavior and dopamine axon activities in visual virtual reality. (**A**) An example scene of the virtual corridor at the starting position. (**B**) (top) The time course of lick rate for each animal (gray) aligned by scene movement onset (*n* = 16 mice) averaged across animals (black). (bottom) Locomotor speed of each animal (gray) averaged across animals (black). The shades represent s.e.m.. Green, red, and blue horizontal bars represent the time windows used to quantify impulsive lick, anticipatory lick, and post-reward lick, respectively, in Fig. 2C. (**C**) (top) Impulsive lick (green), anticipatory lick (red), and post-reward lick (blue) rates as a function of days of training. Asterisks mark days in which lick rates significantly differed from those on the first day of training (*n* = 16, signed rank test, alpha = 0.05). Anticipatory lick increased (*R* = 0.39, *P* = 2.7 × 10^-7^), impulsive lick decreased (*R* = –0.36, *P* = 3.9 × 10^-6^), and post-reward lick did not change (*R* = 0.04, *P* = 0.64) over days of training. (bottom) Locomotor speed in the same format as (top). 10 out of 16 mice showed the average running speed > 5cm/s. Error bars represent s.e.m.. (**D**) Schematic diagram of fiber fluorometry (photometry) experiment for monitoring axonal calcium signals. The virus expressing a calcium indicator (GCaMP6m) in a Cre-dependent manner was injected into VTA and SNc, and optical fibers were implanted in the ventral striatum. Fluorescent signals were collected from dopamine axons in the ventral striatum. (**E**) Recording locations in the experimental (green) and control (red) animals (*n* = 16 and 6 mice, respectively). (**F**) Z-scored dopamine axon calcium signals in the ventral striatum from individual animals (gray, *n* = 16) are averaged (black). A gray horizontal bar depicts a temporal window used to compute Pearson’s correlation between time points and fluorescent signals (Ramping *R*; see Methods). (**G**) Ramping *R*s (Pearson’s correlation coefficients between ΔF/F and time) at different days of training. Dots represent data from individual animals. Error bars represent s.e.m.. Asterisks indicate that the median was significantly different from zero (signed rank test). (**H**) Z-scored GFP signals from dopamine axons in the ventral striatum. (**I**) GFP signals were not significantly different from zero (*P* > 0.05, signed rank test for each day, *n* = 6).

We first performed fiber fluorometry (photometry) (Howe and Dombeck, 2016; Kudo et al., 1992; Lerner et al., 2015; Parker et al., 2016) to measure dopaminergic axon activity in the ventral striatum (Babayan et al., 2018; Menegas et al., 2017, 2018) (Fig. 2D,E). A calcium indicator, GCaMP6m (Chen et al., 2013b), was expressed in dopamine neurons in the VTA and the substantia nigra pars compacta (SNc). The bulk calcium signal was recorded from their axons through an optic fiber implanted in the ventral striatum. We observed that as animals were trained on the task, axonal calcium signals progressively ramped up over the timescale of 3-4 seconds as the animal approached the goal (Fig. 2F,G, Fig. S2E-H). We first quantified the ramping signal in each session using the correlation coefficient (“ramping *R*”) between calcium signals and time (*n* = 16 mice, mean correlation coefficient, *r* = 0.18 ± 0.04; correlation using the average calcium trace was 0.45; 95% confidence interval (CI) = [0.43, 0.48], *P* < 10^-20^; see Methods). Across sessions, anticipatory licking was significantly correlated with the ramping signal while running speed was not (beta for lick = 0.018, 95% CI = [0.0021 0.033]; beta for locomotion speed = 0.0043, CI = [-0.0035 0.012], *n* = 93 sessions, multiple linear regression, see Methods). These ramping dopamine signals are unlikely to be a correlate of motion artifacts or licking, however. First, to examine the potential contributions of motion artifacts, we performed control experiments using mice in which GFP, instead of GCaMP6m, was expressed in dopamine neurons.The result showed that GFP control mice did not exhibit ramping signals (Fig. 2H,I). Second, we have not observed ramping signals in our previous studies using similar techniques (Babayan et al., 2018; Menegas et al., 2017, 2018) in the absence of dynamic sensory stimuli although mice exhibited a similar pattern of anticipatory licking in these conditions.

### Ramping dopamine signals in visual virtual reality are consistent with reward prediction errors

After confirming that we observed a dopamine ramp, we performed a set of 4 experiments designed to explicitly dissociate the two hypotheses. In Experiment 1, in addition to the standard condition, we randomly interleaved three types of ‘test’ trials. Test trials included either a long teleport, a short teleport, or a 5-sec pause (Fig. 3A; **Supplementary Movies 2-4**). The two types of teleport instantaneously moved the animal to the same location (location, 70 a.u.; the start and end of the linear track, 0 and 100 a.u., respectively; the goal location, 97 a.u.), and in ‘pause’ trials, the scene movement was temporarily frozen at this location (70 a.u.). If dopamine represents state value (in this case, the value of locations in the linear track), dopamine signals would be predicted to show a step-like increase arriving at the same level after both a long and short teleport (Fig. 3B left). If dopamine represents RPE, dopamine signals would be predicted to show a phasic (short-lived) excitation, whose magnitude scales with the length of teleport (Fig. 3B right). Value depending on the distance to reward will stay constant when the scene movement is paused, whereas RPE will decrease back to baseline since there is no change of value in time. We note that there is some ambiguity as to how ‘value’ may behave in the pause condition: for example, if the animal judges that the task is aborted at the time of pause, the value may also decrease back to baseline similar to RPEs. We used the results holistically to judge which hypothesis parsimoniously accounts for the entire data in the following.

**Fig. 3.**
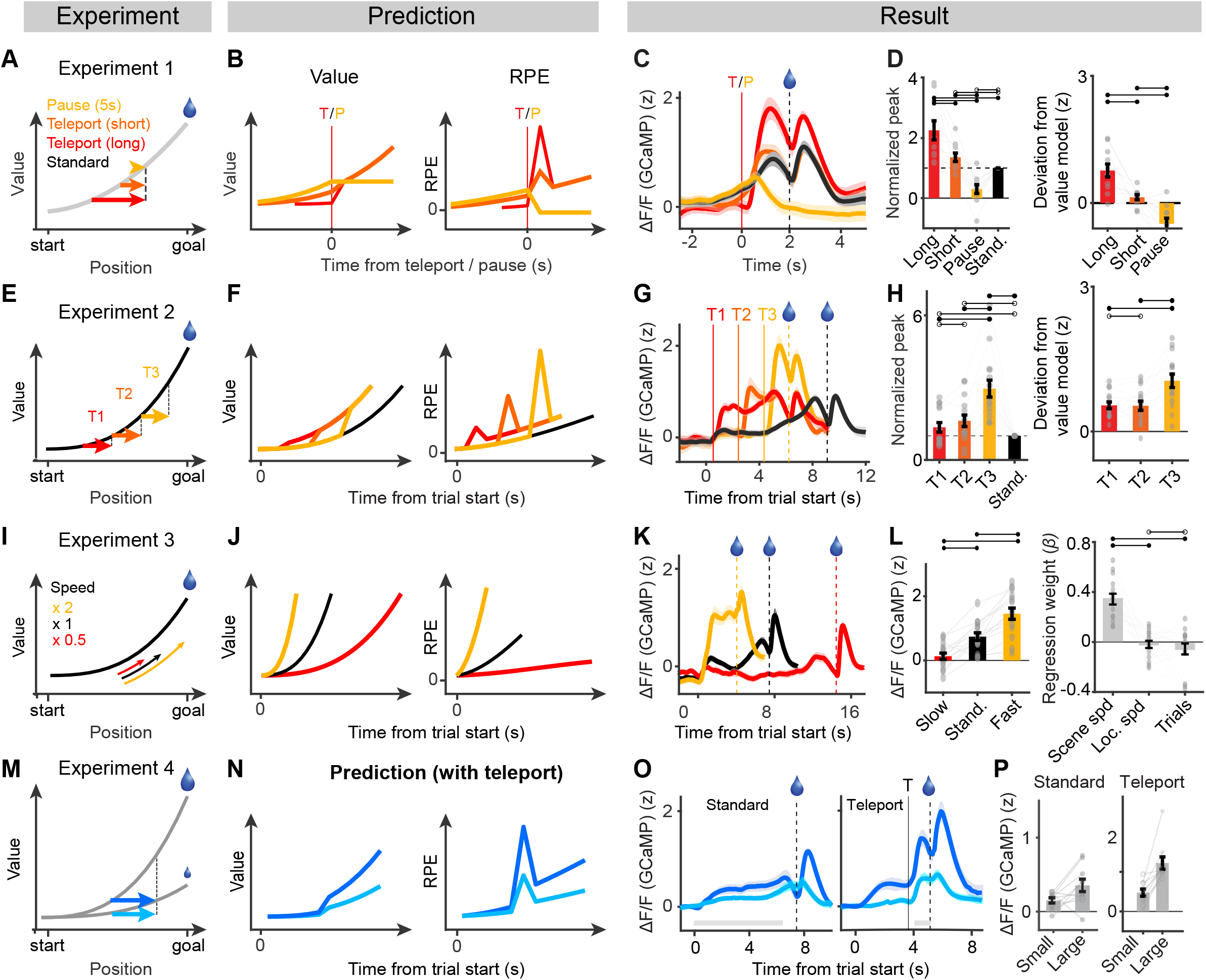
Dopamine axon activities in the ventral striatum are consistent with the RPE hypothesis. (**A**) Experiment 1 design. Long-distance teleport (L, red), short-distance teleport (S, orange), and pause (P, light orange) conditions are depicted on the value function. (**B**) Predictions of how the state value (left) and the RPE (right) change before receiving reward in each experimental condition. T, teleport. P, pause. (**C**) Average calcium signals (z-score) across animals aligned by teleport or pause (*n* = 11 mice). Format as in (B). Shading represents s.e.m.. Calcium signals in the standard condition (black) are aligned such that water delivery time matches with teleport conditions. (**D**) (left) Comparison of peak responses normalized by the peak of ramping in the standard condition (*n* = 11 mice). (right) Residuals from the state value prediction (*n* = 11 mice, see Methods and Fig. S9). Error bars represent s.e.m.. (**E**) Experiment 2 design. Teleports at three different positions (T1, T2, T3) are depicted on the value function. (**F**) Predictions of state value (left) and RPE (right) models. The RPE model predicts that the magnitudes of phasic responses should increase with proximity to the goal. (**G**) Average calcium signals (*n* = 11 mice). (**H**) (left) Normalized peaks increase with proximity to the reward location. (right) Residuals from the state value prediction (median *Test R* = 0.20, *n* = 15 mice, *P* = 0.0031, signed rank test). (**I**) Experiment 3 design. Manipulations of speed depicted on the value function. (**J**) Predictions of state value (left) and RPE (right). The state value should reach the same maximal level at the goal location (left). RPE should increase as a function of scene speed (right). (**K**) Average calcium signals (*n* = 15 mice). (**L**) (left) Comparison of average pre-reward responses at [–1 s 0 s] relative to reward. (right) Regression analysis (see Methods). The pre-reward responses (ramping signals) were regressed on the scene speed, locomotion speed, and trial number. The ramping signals are correlated with the speed of scene movement but not with the animal’s locomotion speed. (**M**) Experiment 4 design. The reward size was altered between small (2.25 µl) and large (10 µl) amounts across blocks of trials. The value function is expected to be scaled according to the reward size. (**N**) Predictions of state value (left) and RPE (right). The value model predicts that step-like changes should be greater in the large-reward block (left). The RPE model predicts that the magnitude of the phasic response should increase in the large-reward block (right). (**O**) Averaged calcium signals with (left) or without (right) teleport (*n* = 10 mice). (**P**) (left) The ramp magnitudes are significantly greater in large-reward blocks than in small-reward blocks (*n* = 10 mice, *P* = 0.049, signed rank test). (right) Phasic teleport responses in large-reward blocks are greater than those in small-reward blocks (*n* = 10, *P* = 0.002, signed rank test).

In teleport trials, anticipatory licking and changes in locomotor speed reflected the new location (Fig. S3B), confirming that mice used the spatial (visual) cues to predict reward rather than merely relying on the elapsed time. A long-distance teleport evoked a large calcium transient whose peak was greater than the peak of the ramp in the standard condition (Fig. 3C, D left, the ratio between the peaks: 2.25 ± 0.31, *P* = 0.0010, *n* = 11 mice, signed rank test; see Fig. S3A for a single session example). The phasic excitation evoked by a short-distance teleport was smaller than that evoked by a long teleport but was still greater than the peak of the ramp in the standard condition (Fig. 3C, D left, ratio: 1.35 ± 0.14, *P* = 0.024). During pause trials, the calcium signal decreased to the baseline level, followed by a phasic excitation when the scene movement resumed (Fig. 3C, D left).

To quantify these results, we generated predicted responses based on the value hypothesis (Fig. S4A-D; see Methods). If dopamine represents value, the deviation from these predictions should be small and unsystematic. If dopamine signals represent RPEs, the deviation from the prediction will be systematically modulated by the conditions: a large positive deviation for long-distance teleport, a small positive deviation for short-distance teleport, and a negative deviation for pause trials. In most of the animals (9 out of 11), the deviations of the measured calcium signals followed this pattern (Fig. 3D right, significance of *Test R*, alpha = 0.05; median *Test R* = – 0.64, *n* = 11, *P* = 0.0020, signed rank test; see Methods for the definition of ‘*Test R*’), supporting the RPE hypothesis.

Since value is a latent variable that cannot be directly observed, the ‘shape’ of the value function cannot be directly assessed. The inability to directly characterize the shape of the value function is a major obstacle in dissociating the value from RPE (Gershman, 2014). In Experiment 2, we sought to infer the shape of the value function using teleport in addition to separating value and RPE (Fig. 3E-H; Fig. S3C, D). In test trials, mice were teleported forward from one of three locations by the same distance (Fig. 3E). If the underlying value function has a convex shape, the magnitude of dopamine response, either in a step-like or transient form, should be larger with teleport occurring at locations closer to the goal. Indeed, the phasic calcium signals followed this pattern (Fig. 3H left, median *r* = 0.45, *P* = 6.1×10^-5^, *n* = 15 mice, signed rank test, Spearman correlation between position and response; Fig. 3H right, median *Test R* = 0.20, *P* = 0.0031, see Methods), supporting the assumption of our experiments — a monotonically increasing value function whose slope increases with proximity to reward (i.e., a convex value function).

The phasic responses at the time of teleport observed in Experiments 1 and 2 demonstrate that dopamine signals approximate the derivative of the value function, consistent with TD RPE.

To obtain further evidence that the ramping itself represents RPEs, we modulated the speed of scene movement (Experiment 3, Fig. 3I-L; Fig. S3E, F; **Supplementary Movies 5,6**). Value depending on the distance to reward would reach the same peak level irrespective of the speed of the scene, whereas the TD RPE signal would increase with scene speed as the speed directly modulates the change of value in time (Fig. 3J). In test trials, we moved the scene either faster (×2 speed) or slower (×0.5 speed) compared to the standard condition. We found that calcium signals reached higher and lower levels in the ‘fast’ and ‘slow’ conditions, respectively, compared to the standard condition (Fig. 3K, L left, *P* = 6.1 × 10^-5^ and 6.1 × 10^-4^, *n* = 15 mice, signed rank test). Regression analysis further showed that the magnitude of ramping can be predicted by the speed of the visual scene, but not correlated with animals’ locomotion speed (Fig. 3L right). These results are incompatible with the value account because dopamine activity should reach the same level at the goal regardless of the speed. Instead, this result can be accounted for by a derivative-like computation, more specifically, the temporal derivative of spatial value (the value associated with different locations), again violating the value hypothesis and supporting the RPE hypothesis.

To examine the potential contributions of motion artifacts, we performed control experiments using mice in which GFP was expressed in dopamine neurons (Fig. S4H-M) in Experiments 1-3. Systematic modulations of signals observed in GCaMP mice (Fig. 3D, H, L) were not observed in these control animals (*n* = 5 mice).

To directly compare which of the value or RPE hypotheses explains the data better, we next performed a model fit analysis. The state value was first defined using a monotonically increasing function over space (Fig. 4A). Based on this state value function, we then predicted calcium signals in each experimental condition according to the value and RPE hypotheses, and obtained a set of parameters that best fit the data (Fig. 4A, B; Fig. S5; see **Methods**). For model comparisons, the goodness-of-fit was quantified using the Akaike information criterion (AIC) to penalize the number of parameters in a model. We first used a value function whose value is discounted by a fixed rate (*r*) as a function of the distance to the target. We found that the RPE model explained the data far better than the value model in all experimental conditions (Fig. 4C, D, *P* < 0.002 for all four fits with manipulated experiments, *n* = 15, 12, 16, 16 for Experiments 1, 2, 3, and all, signed rank test). In the standard condition, however, there was no significant difference in the goodness-of-fit between the RPE and value models (Fig. 4D, ‘Standard’; *P* = 0.49, *n* = 16 mice), indicating that our method is unbiased. The superiority of the RPE over value model was consistent irrespective of the function used to approximate the state value (Fig. 4E). These results indicate that the calcium signals that we observed in Experiments 1-3 are better explained by TD RPEs than by state values.

**Fig. 4.**
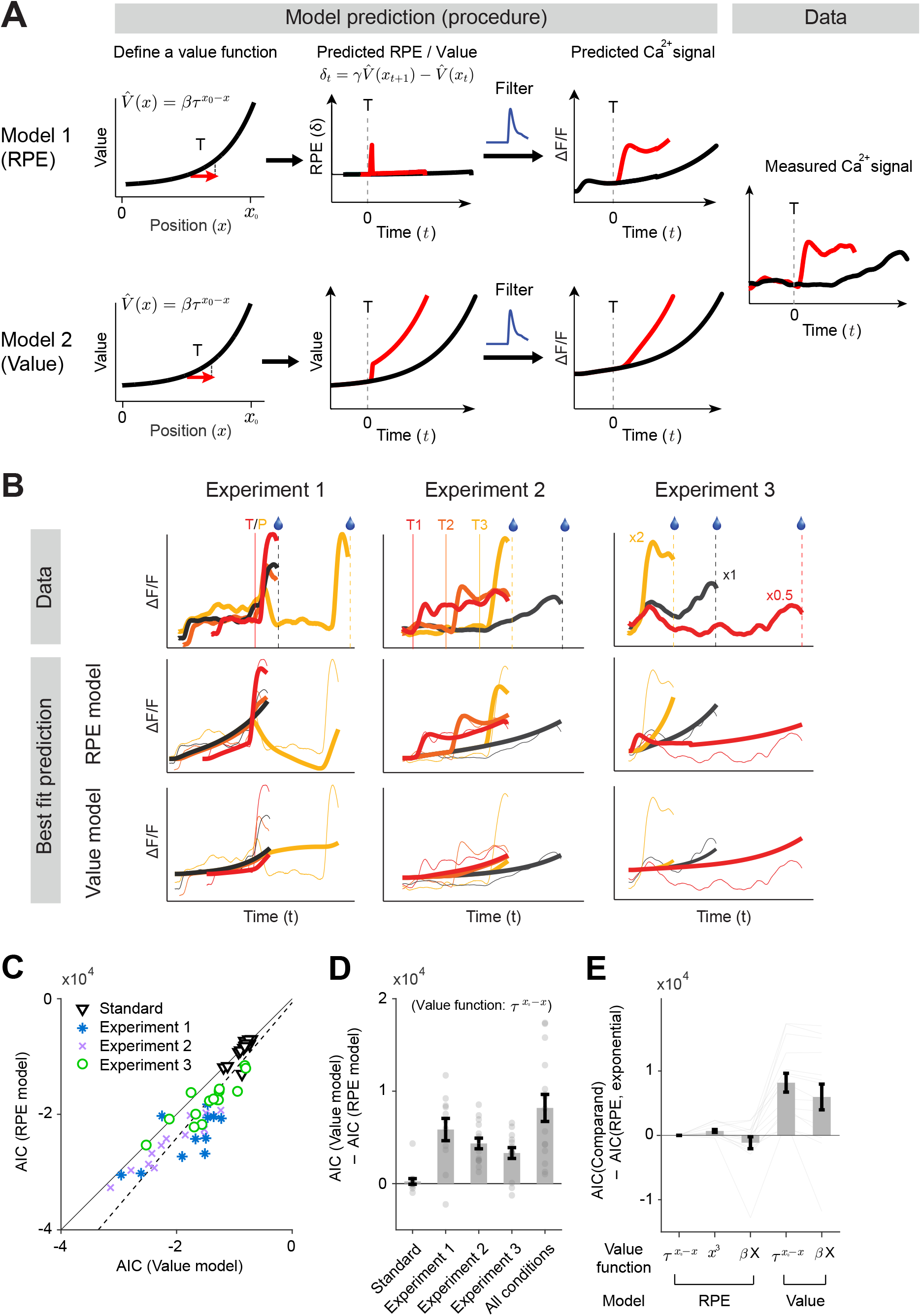
RPE models explain the data better than value models. (**A**) A schematic illustration of the model fitting procedure. Value functions were defined by a set of parameters. Here an exponential decay function (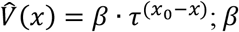, maximum value; *x_0_=* 97, goal location; *τ*, spatial discounting factor) was used for illustration. For TD RPE models, TD RPEs were computed based on the value function with the temporal discounting factor (*γ*) as an additional parameter. The resultant responses (Predicted RPE) were convolved with the GCaMP filter (blue) to generate predicted calcium responses. Non-linear fitting procedure was applied to obtain fitting parameters that minimize the squared sum of errors (See Methods). (**B**) Example fitting results for all conditions. (top) Calcium signals in Experiments 1-3. (middle) Best fit results for the RPE model. One model was fit to all the conditions (Experiments 1-3). Thick lines, model prediction. Thin lines, data (same as the top). (bottom) Best fit results for the value model. Same convention as the middle panels. The RPE and value models were separately fit to the data. (**C**) Comparisons of goodness-of-fits quantified by AIC between value and RPE models based on the exponential value function (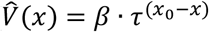, See Methods). (**D**) Difference between the two models. “All conditions” include the data from Experiments 1-3. Error bars represent s.e.m.. (**E**) AICs from variants of models relative to the RPE model in (c), using combined dataset. Different definitions of value function are used. *τ*^(*x*_0_−*x*)^, exponential discounting; *x^3^*, cubic; *βX = β_0_+ β_1_x + β_2_x^2^+ β_3_x^3^*, polynomial (See Methods). Error bars represent s.e.m..

We next examined whether the magnitudes of both ramping and teleport responses are sensitive to reward values as has been previously proposed (Howe et al., 2013). In Experiment 4, the amount of reward was altered across blocks of trials (Fig. 3M-P). In large-reward blocks, mice showed greater anticipatory licking and post-reward licking compared to small-reward blocks (*P* = 0.008, *n* = 10, signed rank test). The magnitude of ramping was greater in large-reward blocks than in small-reward blocks (Fig. 3O left**, 3P** left, *P* = 0.049, *n* = 10 mice, signed rank test) (Howe et al., 2013). Furthermore, the phasic response to teleport was greater in large-reward blocks (Fig. 3O right, **3P** right, *P* = 0.0020, *n* = 10 mice, signed rank test). These results demonstrate that both ramping and transient responses to teleports are sensitive to outcome values and cannot be accounted for by a purely sensory surprise. Using a subset of animals, we also confirmed that dopamine activity was not modulated by a purely sensory aspect of teleport (discontinuity of visual scene) or speed of motion when the animals were not trained on the task with reward (see Methods for ‘No reward control experiments’; Fig. S4E-G).

Together, the results from Experiments 1-4 strongly contradict the hypothesis that dopamine ramps represent state values. Instead, they can be parsimoniously explained by the TD RPE hypothesis—a derivative-like computation over a convex value function (more precisely, a “sufficiently” convex function, as detailed in **Supplemental Note**; Fig. S1).

### Ramping signals originate in the spiking activity in the VTA

The above results indicate that the activity of dopamine axons in the ventral striatum conforms to various predictions of RPEs. However, it remains unclear whether these results hold at the single neuron level. For instance, it is important to determine whether a single neuron exhibits both ramping and teleport-induced transient responses as the RPE hypothesis predicts. Furthermore, a recent study concluded that the spiking activity of VTA dopamine neurons does not ramp up, and ramping dopamine signals arise due to local modulations at their axons independent of the activity at cell bodies (Mohebi et al., 2019). To address these issues, we next characterized the spiking activity of dopamine neurons in mice performing our tasks. We recorded from the lateral VTA that is enriched by dopamine neurons projecting to the ventral striatum. To unambiguously identify dopamine neurons, the light-gated cation channel channelrhodopsin-2 (ChR2) was expressed in dopamine neurons. We classified the recorded neurons as dopaminergic when they responded to light reliably with short latency (19 neurons in 4 animals) (Fig. 5A-D) (Cohen et al., 2012; Eshel et al., 2015; Lima et al., 2009).

In the standard condition, most of the optogenetically identified dopamine neurons showed a strong phasic excitation in response to reward (Fig. 5E, *P* = 0.004, *n* = 19 neurons, signed rank test; 16/19 neurons show significant excitation, alpha = 0.05, signed rank test) consistent with previous studies (Cohen et al., 2012; Eshel et al., 2016; Tian and Uchida, 2015). In addition, the firing rate of some dopamine neurons increased or decreased before receiving reward with some dopamine neurons gradually ramping up or ramping down (Fig. 5G top). On average, the spiking activity exhibited a moderate ramping, increasing by 1.89 ± 0.44 spikes per second from baseline to the peak of the ramping (bootstrap with 1000 repetitions; median ramping *R*, *r* = 0.05, *P* = 0.006, signed rank test. *n* = 19 neurons). Since the magnitude of ramping was markedly smaller compared to the magnitude of reward response, we asked whether the small increase in firing rate can sufficiently account for the ramping signals measured based on the calcium signals described above (Fig. 2, 3). We reasoned that the key difference may lie in the different kinetics between spike and calcium signals. That is, a single spike produces a seconds-timescale increase in calcium due to the slow dynamics of calcium and the kinetics of calcium indicators (including GCaMP6m that we used). To test our hypothesis, we generated predicted calcium signals based on the spiking activity obtained in our recording experiment (see Methods). We found that predicted calcium signals made the ramping signals more prominent and the phasic responses less prominent, compared to the raw firing rates (Fig. 5F). Thus, the slow kinetics of calcium signals exaggerate the slow-timescale changes in spikes while deemphasizing phasic spike responses. Taken together, the spiking activity of VTA dopamine neurons exhibit a positive ramp as a population, which can account for the ramping calcium signals observed in the above experiments.

**Fig. 5.**
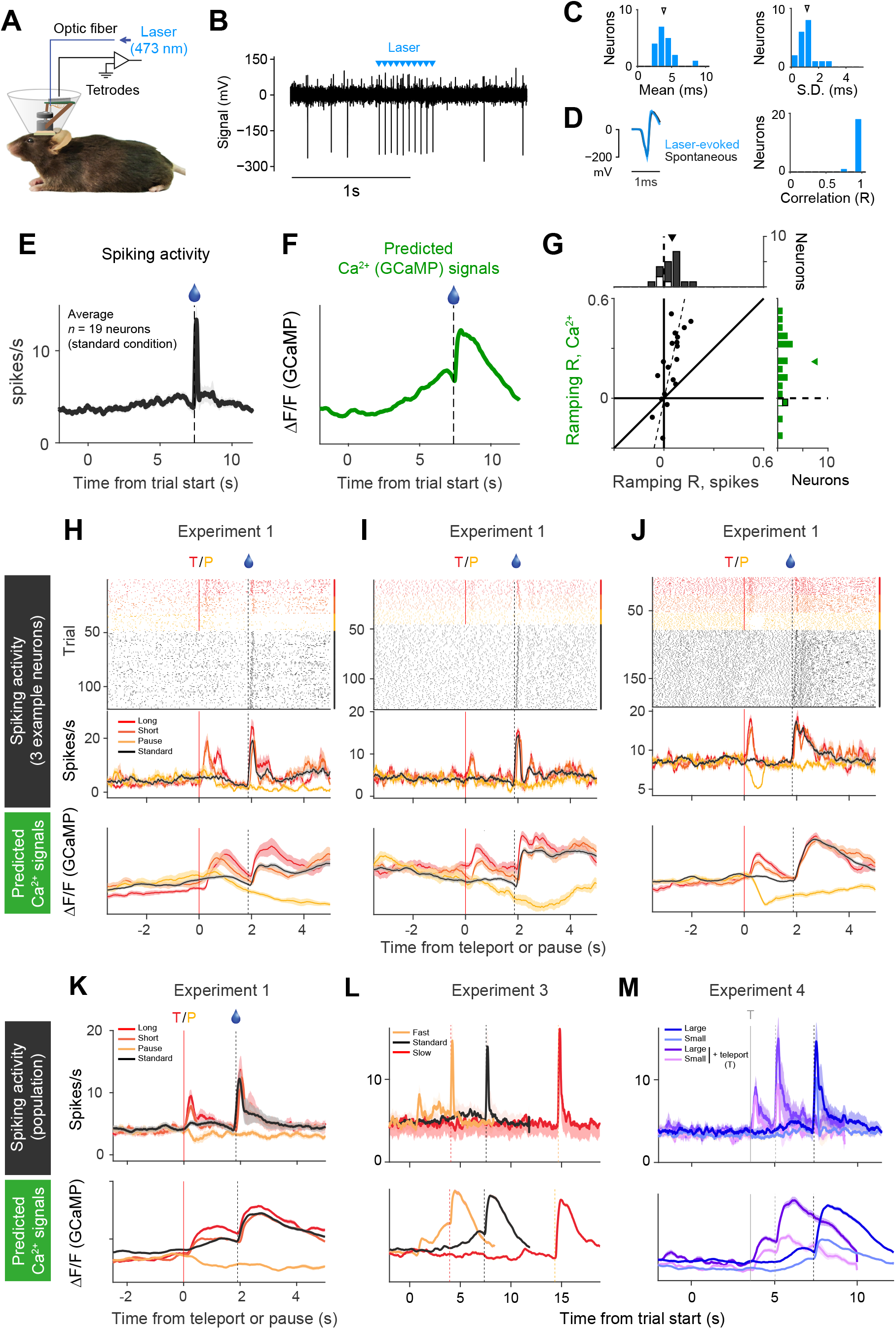
Spiking activity of dopamine neurons in VTA can account for the ramping calcium signals. (**A**) Illustration of chronic tetrode implantation with an optic fiber (**B**) The raw spike recording data during optogenetic identification of dopamine neurons. This neuron reliably responded to laser pulses (cyan, 20Hz, 5ms) (**C**) Latency of laser-evoked spikes. (left) Histogram of average latencies across neurons (*n* = 19). Triangle denotes the median (3.85 ms). (right) Histogram of the standard deviations of the average latencies across neurons (*n* = 19). Triangle denotes the median (1.16 ms). (**D**) (left) Comparison of spike waveforms between laser-evoked spikes (cyan) and spontaneous spikes (black). (right). Histogram of correlation coefficients between spike waveforms between laser-evoked and spontaneous spikes. All of the identified neurons were significantly modulated by laser pulses (*P* < 0.05, SALT test) (Kvitsiani et al., 2013). (**E**) Average time course of spiking activity in the standard condition of the linear track task. (**F**) Calcium signals predicted from spiking activities. We convolved spikes with a GCaMP kernel, and the results were pooled across neurons (see Methods). (**G**) (top) Histogram of Ramping *R*s based on firing rates. The median of the distribution (triangle, 0.05) is significantly greater than zero (*n* = 19, signed rank test). (bottom right) Histogram of Ramping *R*s based on convolved responses of individual neurons. Median (triangle, 0.22) is greater than zero (*n* = 19, signed rank test). (bottom left) Type 2 regressions show that the GCaMP kernel significantly amplified Ramping *R*s based on firing rates (dotted line, regression coefficient = 4.9, CI = [3.1 8.9]). (**H-J**) Spiking activity and predicted GCaMP signals of 3 example neurons in Experiment 1. (**H**) An example neuron that showed positive ramping (Ramping *R*, *r* = 0.024, *P* < 10^-14^). This neuron showed a phasic excitation at the time of teleport. (**I**) A neuron with negative ramp (Ramping *R*, *r* = –0.0214, *P* = 0.0001). (**J**) A neuron without clear ramping. Notably, all three neurons show clear phasic responses to long-distance teleport, which is larger than short-distance teleport. (**K-M**) (top) Averaged spiking activity across population (*n* = 18) in Experiment 1 (**K**), Experiment 3 (*n* = 17) (**L**), Experiment 4 (*n* = 11) (**M).** (bottom) Predicted GCaMP signals based on the convolved responses from the same population of neurons (see Methods for details).

We next examined whether single neuron activities are consistent with RPE or state value using Experiments 1, 2, and 4. We found that a large fraction of neurons, including those that have a positive or negative ramp in the standard condition, exhibited phasic responses at the time of teleport (14/18 neurons significantly greater than the response in the standard condition, *P* < 0.05, signed rank test). Importantly, neurons showed phasic responses to teleport irrespective of whether their overall activity ramped up or ramped down (*R* = –0.15, *P* = 0.55; *n* = 18, Spearman correlation between response to the long teleport and Ramping *R*). Furthermore, the magnitude of ramping was modulated by the velocity of the scene movement (median Test *R* = 0.18, *P* = 0.0003; 7/17 neurons significantly different from zero). According to Gershman (2014), RPEs may ramp up or down depending on the shape (convexity) of the value function that each dopamine neuron uses to compute RPEs (Fig. S1). Furthermore, as discussed above, transient responses at the teleport is a key prediction of TD RPE that can be explained as the temporal derivative of value function as the value suddenly increases at the teleport. These results indicate that although single neurons differ in terms of whether they ramp up or ramp down, the activities of most single neurons are consistent with RPEs.

The above results indicate that somewhat larger ramping signals in axonal calcium signals are due to the slow kinetics of calcium signals compared to spiking activities. Furthermore, the spiking activity at the cell body is sufficient to explain the axonal activity in the ventral striatum. If these were true, then calcium signals measured at the cell bodies of dopamine neurons should also exhibit a similar level of ramping as those at the axons. To test this prediction, we next measured calcium signals in dopamine neurons in the VTA using fiber fluorometry (Fig. 6A; Fig. S6). Indeed, we found that the calcium signals measured through an optical fiber implanted in the VTA showed a similar level of ramping as well as relatively smaller phasic excitation in response to reward. Further testing in Experiments 1-3 confirmed that the main features reported above were evident from the VTA calcium signals and show a striking similarity to the predicted calcium dynamics from the population of VTA neurons (Fig. 5K-M bottom panels correspond to Fig. 6B-D).

**Fig. 6.**
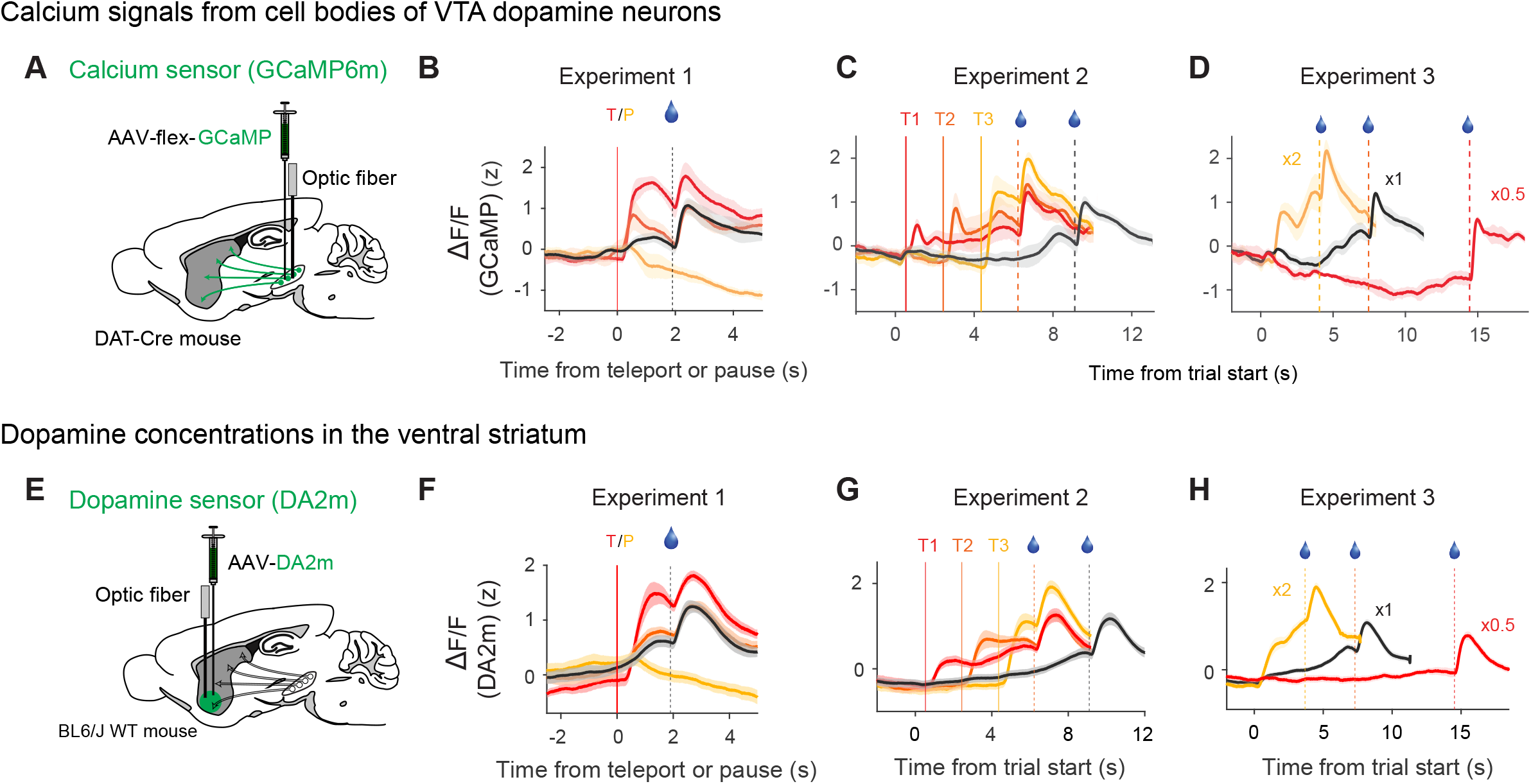
Calcium signals in cell bodies of VTA dopamine neurons as well as dopamine concentrations in the ventral striatum signal RPEs. (**A**) Schematic diagram of fiber fluorometry surgery for GCaMP recording from dopamine neurons in VTA. An optical fiber was implanted into VTA. (**B-D**) Averaged calcium signals in Experiment 1 (B, *n* = 4), Experiment 2 (*n* = 4), Experiment 3 (*n* = 3). (**E**) Schematic diagram of fiber fluorometry surgery for dopamine sensor. A dopamine sensor (GRAB_DA2m_) was expressed in the ventral striatum. An optical fiber was implanted into the ventral striatum. (**F-H**) Averaged dopamine signals in Experiment 1 (**F**, *n* = 9), Experiment 2 (**G**, *n* = 10) and Experiment 3 (**H**, *n* = 10).

### The dynamics of dopamine concentration in the ventral striatum is consistent with reward prediction errors

As discussed above, TD RPE is approximately the derivative of value. This indicates that if the RPE signal is temporally integrated, the RPE signal could be converted to a value-like quantity. The above result showed that although calcium signals are a temporally integrated version of the spiking activity at a 1-2 second timescale, they were still consistent with RPEs (Experiments 1-4). What about the dopamine signals in the projection site? This is not an obvious question because the time course of dopamine can be affected by several factors including dopamine reuptake activity and terminal modulation of dopamine release.

To address this question, we next turned to direct measurements of dopamine using a recently developed genetically encoded dopamine sensor, GRAB_DA_, an improved version based on GRAB_DA1m_ (Sun et al., 2018), with additional point mutations in the inserted circularly permuted EGFP. A dopamine sensor (DA2m) was expressed in neurons in the ventral striatum, and fluorescent signals were measured through an optical fiber implanted in the ventral striatum (Fig. 6E). We then monitored the dynamics of dopamine signals while mice were tested in Experiments 1-3. We found that the dopamine signals that we obtained are very similar to the calcium signals in other experiments (Fig. 6F-H). Importantly, transient excitation to long-distance teleport exceeded the peak of ramping in the standard condition (Fig. 6F, *P* = 0.004, *n* = 9, signed rank test), and the magnitude of ramping signals was significantly different depending on the velocity of scene movement (Fig. 6H, median Test *R* = 0.43, *P* = 0.002, *n* = 10, signed rank test; 9/10 animals showed significant Test *R*, alpha = 0.05). These results are inconsistent with the possibility that the concentration of dopamine represents state values.

### A moving bar that indicates the proximity to reward causes dopamine ramps consistent with reward prediction errors

Although we observed dopamine ramps in the present study, many other studies using classical conditioning have not typically observed dopamine ramps regardless of the techniques used such as electrophysiology (Cohen et al., 2012; Schultz et al., 1997), fiber fluorometry (Babayan et al., 2018; Menegas et al., 2017), and voltammetry (Stuber et al., 2008; Flagel et al., 2011; Hart et al., 2014). Furthermore, ramping signals were not observed regardless of recording locations including cell bodies in the VTA (Cohen et al., 2012; Eshel et al., 2015; Schultz et al., 1997), axonal activities (Menegas et al., 2015) in the ventral striatum or dopamine release (Flagel et al., 2011; Hart et al., 2014; Stuber et al., 2008) in the ventral striatum. Based on these previous observations and the aforementioned results in spike recording, fiber fluorometry of calcium signals in VTA and fiber fluorometry of dopamine concentration in the ventral striatum, the presence or absence of ramping signals does not appear to be due to the techniques used or recording locations, but is likely due to the difference in task conditions.

In previous studies in which dopamine ramps were observed, animals typically traversed or navigated across space to obtain reward (Hamid et al., 2016; Howe et al., 2013; Mohebi et al., 2019; Phillips et al., 2003; Roitman et al., 2004). These conditions differ in at least two ways from the conditions in which dopamine ramps were not observed. First, animals are engaged in voluntary locomotor movement. Second, while moving, the visual input can provide continuously updated sensory information indicating the proximity to reward, which is absent in typical classical conditioning paradigms in which the timing of reward is defined by an elapsed time. In our virtual reality task, the animal was not required to move, although some animals displayed locomotor activities (Fig. 2B bottom). Nonetheless, locomotor activities and the magnitude of ramping signals were not correlated as discussed above (Fig. 3L right). We therefore hypothesized that a sensory input that signals a gradual approach to reward could underlie a dopamine ramp. We tested this idea using a more abstract, non-navigational stimulus: a black horizontal bar that moved downward from the top of computer monitors. The animal received a drop of water when the bar reached a certain location (Fig. 7A, see Methods). Once the animal learned the task (Fig. S7A), we observed a dopamine ramp similar to that in the linear track task (Fig. S7B, C). Note that this task is very similar to classical conditioning tasks used in previous studies except for the presence of a dynamic stimulus.

**Fig. 7.**
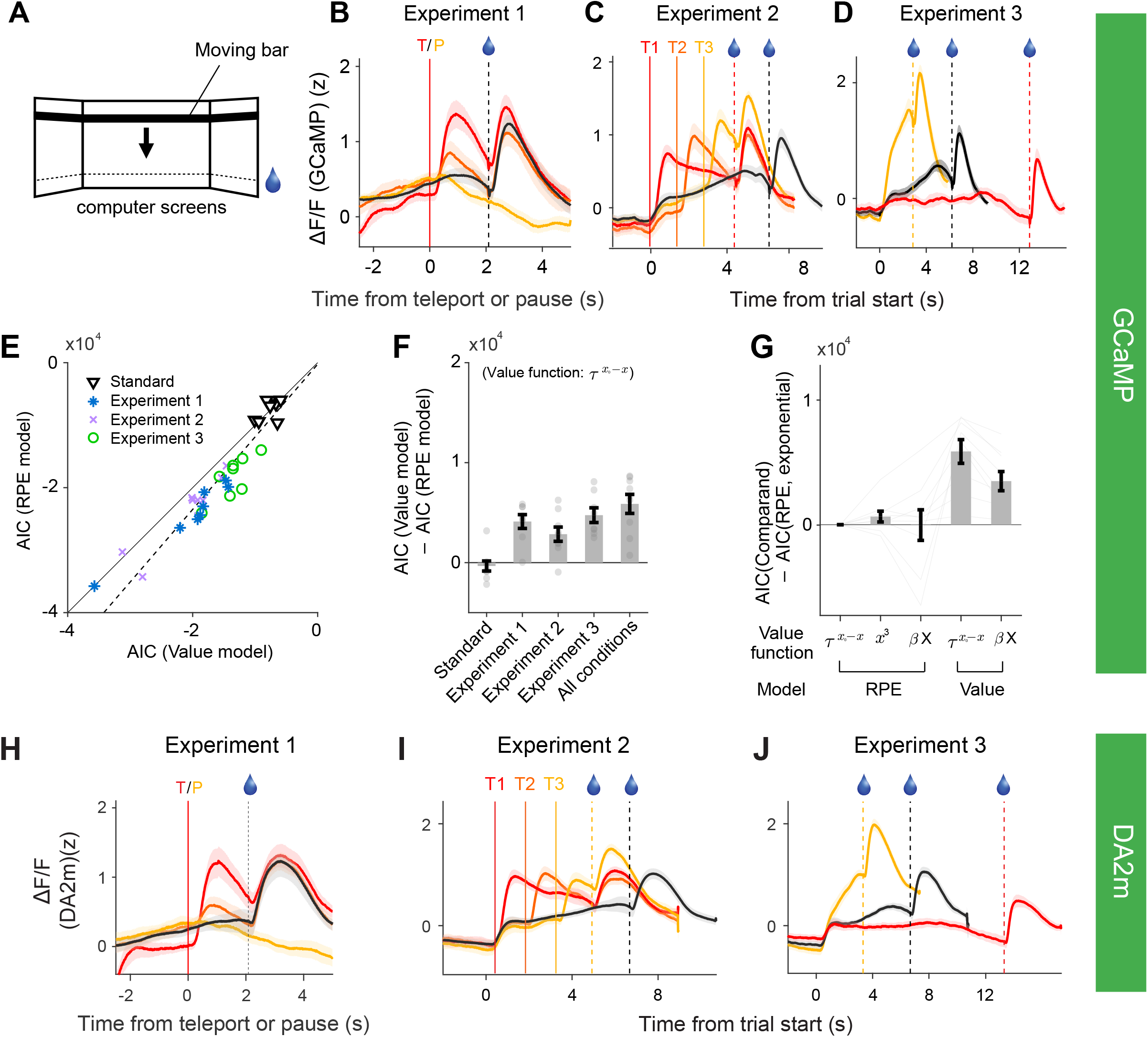
Dynamic sensory stimulus indicating reward proximity can cause a dopamine ramp in a manner consistent with the RPE hypothesis. (**A**) Experimental design. Illustration of the moving-bar experiment. A horizontal bar displayed on the three surrounding screens moved downward at a constant speed. A drop of water was delivered when the bar reached a target position (dotted line, illustration purpose only). (**B**) Experiment 1. Averaged calcium signals across animals aligned by long-distance bar teleport (red), short-distance bar teleport (orange), and pause (light orange) (*n* = 9 mice). Calcium signals in the standard condition (black) are aligned such that water delivery time (black dash) matches with the teleport conditions. Shading represents s.e.m.. (**C**) Experiment 2. Averaged calcium signals to three teleports aligned by bar movement onset (*n* = 10 mice). Vertical lines indicate teleports at high (0.4 s, light orange), middle (1.37 s, orange), and low (2.78 s, red) positions. Red and black dashes depict water delivery for the test and standard conditions, respectively. (**D**) Experiment 3. Averaged calcium signals in the speed manipulation task (*n* = 9 mice). Responses in the ×2 (light orange), standard (×1, black), and ×0.5 (red) conditions are aligned by bar movement onset. (**E**) Comparisons of goodness-of-fits quantified by AIC between value and RPE models based on the exponential value function 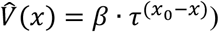. (**F**) Difference between the two models. “All conditions” include the data from Experiments 1-3. All median ΔAICs except for the standard condition are significantly different from zero (*P* = [0.20 0.008 0.02 0.04 0.004], *n* = [9 8 8 9 9], signed rank test). Error bars represent s.e.m.. (**G**) AICs from variants of models relative to the RPE model in (e), using a combined dataset. Different definitions of value function are used. *τ*^(*x*_0_-*x*)^, exponential discounting; *x^3^*, cubic; *βX = β*_0_*+ β*_1_*x + β*_2_*x*^2^*+ β*_3_*x*^3^, polynomial (See Methods). (**H-J**) Averaged dopamine signals measured using a dopamine sensor (GRAB_DA2m_). (**H**) Experiment 1, (**I**) Experiment 2, and (**J)** Experiment 3. The results were similar to those obtained using a calcium sensor (**B-D**).

To evaluate whether the ramping caused by the moving bar was due to an RPE or value signal, we next performed a series of experiments analogous to Experiments 1-3 (see Methods). The results obtained in these experiments were largely consistent with those obtained using the navigational scene stimuli. We observed phasic excitations at the time of bar teleport whose size increased with the teleport distance, and a reduction of signals at pause (Fig. 7B). The peak of the phasic response at a long teleport was significantly greater than the peak of ramping in the standard condition (Fig. S7D, G, *P* = 0.039, *n* = 9 mice, signed rank test). The teleported bar at three different positions evoked phasic responses (Fig. 7C). The magnitudes of the responses showed an increasing trend with teleport positions, although it did not reach statistical significance (Fig. S7E, H, *Test R* = 0.21, *P* = 0.26). Finally, the speed of bar movement strongly modulated the ramping signals (Fig. 7D). The ramp and the speed of bar movement were positively correlated in all animals (Fig. S7F, I, 11/11 mice, median *Test R* = 0.38, *P* = 0.0010, signed rank test). The TD RPE model explained the data better than the value model across animals and experimental conditions, as judged based on a model fit analysis (Fig. 7E-G). Furthermore, dopamine signals in the ventral striatum, measured using GRAB_DA_ sensor (DA2m), showed very similar results as those based on calcium signals (Fig. 7H-J). Together, these results demonstrate that dynamic sensory cues that indicate a gradual increase in the proximity to reward can cause a dopamine ramp in a manner consistent with TD RPEs.

## DISCUSSION

In the present study, we examined the nature and the origin of ramping dopamine signals. We developed a set of experimental paradigms to dissociate the two alternative hypotheses (RPE versus value) that are otherwise difficult to disambiguate. Importantly, these experiments are designed based on the original mathematical definition of TD RPE – a derivative-like computation over value (Doya, 2000; Sutton, 1988; Sutton and Barto, 1990, 1998). Using these paradigms, we found that the manipulation of scene movement – teleport and speed manipulations – caused ramping and phasic dopamine responses in the ventral striatum that were consistent with TD RPEs but inconsistent with state values. Furthermore, this was true at all of the stages of dopamine transmission that we examined – spiking activities and calcium signals at cell bodies, calcium signals at axons, and dopamine concentrations in the ventral striatum.

Additionally, we found that a more abstract, non-navigational stimulus that indicates temporal proximity to reward is sufficient to cause a dopamine ramp. These results indicate that the mere presence of ramping dopamine signals does not indicate that they represent values, and ramping dopamine signals in the ventral striatum represent TD RPEs, at least in our experimental conditions. To our knowledge, this is the first experimental demonstration that slowly-fluctuating dopamine signals at a seconds-timescale also encode RPEs, similar to phasic dopamine signals. Together, our results provide a unified account of dopamine signals across these timescales as RPEs.

A derivative-like computation has been at the heart of the TD RPE since its birth (Sutton, 1988; Sutton and Barto, 1990). However, it has not been tested experimentally whether dopamine neurons compute the derivative of value on a moment-by-moment basis when there are no discrete, salient events. We find that dopamine signals in the ventral striatum can be parsimoniously explained as the temporal derivative of the convex value function that is associated with spatial location. The results thus uncover the “mathematical formula” of dopamine RPE signals in our tasks in unprecedented fidelity. Together, our results indicate that the RPE account of dopamine responses can be extended to slowly fluctuating dopamine signals in addition to phasic dopamine responses, and support the previously untested central tenet of TD RPEs that dopamine neurons signal RPEs through a derivative-like computation over value on a moment-by-moment basis.

### The origin of ramping dopamine signals and the potential role of state uncertainty

In the present study, we studied the origin of ramping dopamine signals by characterizing various stages involved in dopamine transmission: (1) spiking activity at the soma using electrophysiological recording, (2) calcium signals at the soma and the axons measured using a calcium indicator (GCaMP6), and (3) dopamine concentrations at the projection site measured using a recently developed dopamine sensor (GRAB_DA2m_). Our results show that ramping dopamine signals observed in the ventral striatum can be readily explained by the spiking activity of dopamine neurons. Although our results do not exclude the possibility that axonal modulations play a role in sculpting dopamine signals in general (Cachope et al., 2012; Sulzer et al., 2016; Threlfell et al., 2012; Zhou et al., 2001), this process is not essential in explaining ramping dopamine signals at least in our paradigms.

The present results suggest that cues that indicate a gradual increase in proximity to reward play a key role in generating ramping dopamine signals. What then explains the difference between tasks in which dopamine ramps up or not? Do the shapes of the value function differ between the tasks? In reinforcement learning, values are computed based on the current “state” of the world. In this framework, the state is collectively defined by various types of information such as the location, the objects that are present there, and the elapsed time from salient events. In natural situations, the state is often ambiguous due to partial information that the animal receives or the internal noise in the brain, and state uncertainty can alter the shape of value functions (Gershman et al., 2014; Ludvig et al., 2008; Starkweather et al., 2017). Importantly, the tasks that we used in the present study (the virtual reality and moving-bar tasks) are different in terms of the structure of state uncertainty from the delayed reward tasks that have been commonly used to characterize dopamine activities. In the virtual reality and moving-bar tasks, sensory inputs continuously provide updated information about the proximity to reward on a moment-by-moment basis, indicating that the uncertainty about the current location (state) relative to the goal location decreases or, at least, stays constant as the distance to the reward location decreases. In contrast, in the absence of such cues, it is known that the uncertainty about the proximity to reward increases proportionally with the elapsed time. That is, in a delayed-reward task, state uncertainty grows as time elapses following Weber’s law, the property called scalar timing (Gibbon et al., 1997). This difference in the structures of state uncertainty may result in different shapes of value functions between the tasks: large state uncertainty in a delayed reward task may dampen the state value closer to the reward location, thus making the value function less convex in shape. We further examine these ideas in the accompanying paper (Mikhael et al.). It will be of interest to directly test these ideas further theoretically as well as experimentally in the future.

### Dopamine signals and behavioral regulation

What dopamine represents and how specific information conveyed by dopamine regulates distinct aspects of behavior remain highly debated. Mohebi et al. (2019) indicated that dopamine concentrations in the ventral striatum are largely independent of the spiking activity of VTA dopamine neurons. Based on our results, we suggest that it is important to divide these results into three timescales: hundreds of milliseconds (often referred to as “phasic”), seconds, and minutes timescales. Our results raise the possibility that seconds-timescale fluctuations of dopamine in Mohebi et al. (2019) could be explained by spiking activities of dopamine neurons. Indeed, after taking into account the slow kinetics of dopamine signals, the spiking data obtained in Mohebi et al. (2019) can readily explain the dopamine sensor signals in their study. These results suggest that a large discrepancy between spiking activities and dopamine release is limited to the minutes-timescale, often referred to as ‘tonic’ dopamine (Niv et al., 2007; Schultz, 2007). It should also be noted that the argument in Mohebi et al. (2019) assumed a linear relationship between spike counts and dopamine release. Whether some non-linear relationship between spikes and dopamine release (e.g. a sigmoidal or saturating function) alone can explain the discrepancy or whether axonal modulation of dopamine release plays an essential role on the minuites timescale remains to be examined.

The present study focused on the question of “representation”, i.e., what information do dopamine neurons convey to the downstream neurons: are they RPE or value? Our results leave open how dopamine signals observed in our paradigm function. Previous studies proposed a dichotomy between slow time-scale, tonic dopamine signals that regulate motivation and phasic dopamine signals that are specialized for reinforcement learning (Berke, 2018; Niv et al., 2007; Schultz, 2007). However, the relationship between specific dopamine signals and different aspects of behavior remains to be established. As discussed above, RPEs can, in principle, be converted to the expected value as the latter is the integral of the former. Although our results showed that dopamine concentrations in the ventral striatum are still closer to RPEs than values, it is conceivable that intracellular or circuit properties downstream of dopamine can temporally integrate RPE signals, thereby computing expected values. In this sense, dopamine RPE signals can, in principle, regulate motivation in a value-dependent manner. Regardless of what dopamine represents, experimental results have indicated that dopamine can modulate both plasticity and excitability of dopamine-recipient neurons (Tritsch and Sabatini, 2012). It is the task of future studies to clarify how distinct functions of dopamine such as motivation and learning arise from distinct types of information conveyed by dopamine.

### Hierarchy in the diversity of dopamine signals

Increasing evidence indicates the existence of diverse dopamine signals. In particular, recent studies have shown that different regions of the striatum receive distinct dopamine signals (Howe et al., 2013; Lerner et al., 2015; Parker et al., 2016; Howe and Dombeck, 2016; Menegas et al., 2017, 2018). Some populations of dopamine neurons in the VTA and SNc are not activated by reward but predominantly by external threats (Menegas et al., 2018), salience (Lerner et al., 2015; Matsumoto and Hikosaka, 2009) or movements (Howe and Dombeck, 2016; Lee et al., 2019; Parker et al., 2016; da Silva et al., 2018). These studies have indicated that these dopamine signals are confined to a subpopulation of dopamine neurons or dopamine signals in specific regions of the striatum.

A recent study showed that single neuron activities of VTA dopamine neurons can be classified into several groups, where each of which is modulated preferentially by specific variables such as “position” (corresponding to the ramping dopamine signals observed here), “speed”, or accuracy (Engelhard et al., 2019). Our results indicate that dopamine signals that are correlated with position and speed (Engelhard et al., 2019) can naturally arise as TD RPE signals. Furthermore, consistent with Engelhard et al. (2019), we observed dopamine neurons with positive, no, or negative ramping activities at the single-neuron level. Importantly, the activity of dopamine neurons with distinct ramping activities also conformed to the predictions of TD RPEs, exhibiting transient activations at the time of teleport and different magnitudes with speed manipulations. Applying the theory proposed by Gershman (2014), these types of variability can be explained by different shapes of value functions (i.e., more or less convex value functions) that underlie RPE computations (Fig. S1). Notably, most dopamine neurons in Engelhard et al. (2019) exhibited typical RPE responses in a delayed-reward task. Taken all together, these results raise the possibility that at least some fraction of variable dopamine responses observed in Engelhard et al. (2019) can be understood as variants of RPEs due to variability in the corresponding value estimators that each dopamine neuron uses to compute RPEs.

Together with other evidence (Watabe-Uchida et al., 2017; Parker et al., 2016; Howe and Dombeck, 2016; Menegas et al., 2017, 2018; Wenzel et al., 2015; but see Yang et al., 2018), the present results reinforce the idea that dopamine signals in the ventral striatum resemble TD RPEs. Although dopamine neurons in this population exhibit some diversity (Engelhard et al., 2019; Yves et al., 2018), at least some of the observed variability can be understood within the framework of RPEs. In contrast, more globally, a much larger level of diversity exists across different regions of the striatum: ventral, dorsomedial, dorsolateral, and the tail of striatum (Cox and Witten, 2019; Watabe-Uchida and Uchida, 2019). These results indicate that the diversity of dopamine neurons is hierarchically organized. While distinct dopamine subsystems can be defined by gross differences in dopamine signals observed across different striatal regions, there is relatively minor diversity in dopamine signals that exists within each subsystem. Looking at the diversity of dopamine signals in this hierarchical manner, it will be important to clarify (1) the computational principle by which each dopamine subsystem operates, and (2) whether relatively minor variations in dopamine signals within each subsystem can be regarded as just “noise” in the system or whether there is any functional significance. Our experimental paradigms, which are based on predictions of computational models, will be a powerful means with which to examine the nature of dopamine signals. A deeper understanding of dopamine signals will aid in addressing these questions in the future.

## ACKNOWLEDGMENTS

We thank Dr. Christopher Harvey for the assistance in setting up virtual reality experiments, and Dr. Dmitriy Aronov for developing VirMEn software. We thank Drs. Benedicte Babayan, William Menegas, and Edward Soucy for developing in-house fiber fluorometry setups. We thank Aleeza Shakeel for the assistance on histology. We thank Dr. John Assad for his critical reading of the manuscript. We also thank the members of the Uchida lab for helpful discussions and critical reading of the manuscript. This work was supported by the NIH grants U19 NS113201 (to N.U. and S.J.G.), R01MH095953 (to N.U.), R01MH101207 (to N.U.), NS108740 (to N.U.), T32GM007753 (to J.G.M.), T32MH020017 (to J.G.M), U01NS103558 (to Y.L.); a Harvard Brain Science Initiative seed grant (to N.U.); the Simons Collaboration on Global Brain (to N.U.); a Harvard Mind Brain and Behavior faculty grant (to S.J.G. and N.U.), a research fellowship from the Alfred P. Sloan Foundation (S.J.G.); and Junior Thousand Talents Program of China (to Y.L.).

## AUTHOR CONTRIBUTIONS

H.R.K. and N.U. conceived and designed the experiments. H.R.K. conducted fiber fluorometry experiments using GCaMP and was involved in all the other experiments. A.M. conducted electrophysiology experiments. P.B. conducted fiber fluorometry experiments using the dopamine sensor. I.T-K. and M.W-U. tested the dopamine sensor in the striatum and advised on using it. F.S., Y.Z., and Y.L. developed the dopamine sensor. J.G.M. and S.J.G. provided theoretical analysis and advised data analysis. H.R.K. analyzed the data and wrote the first draft and all the other authors discussed the results with H.R.K. and commented on the manuscript.

## DECLRATATION OF INTERESTS

The authors declare no competing interests.

---

**Fig. S1.**
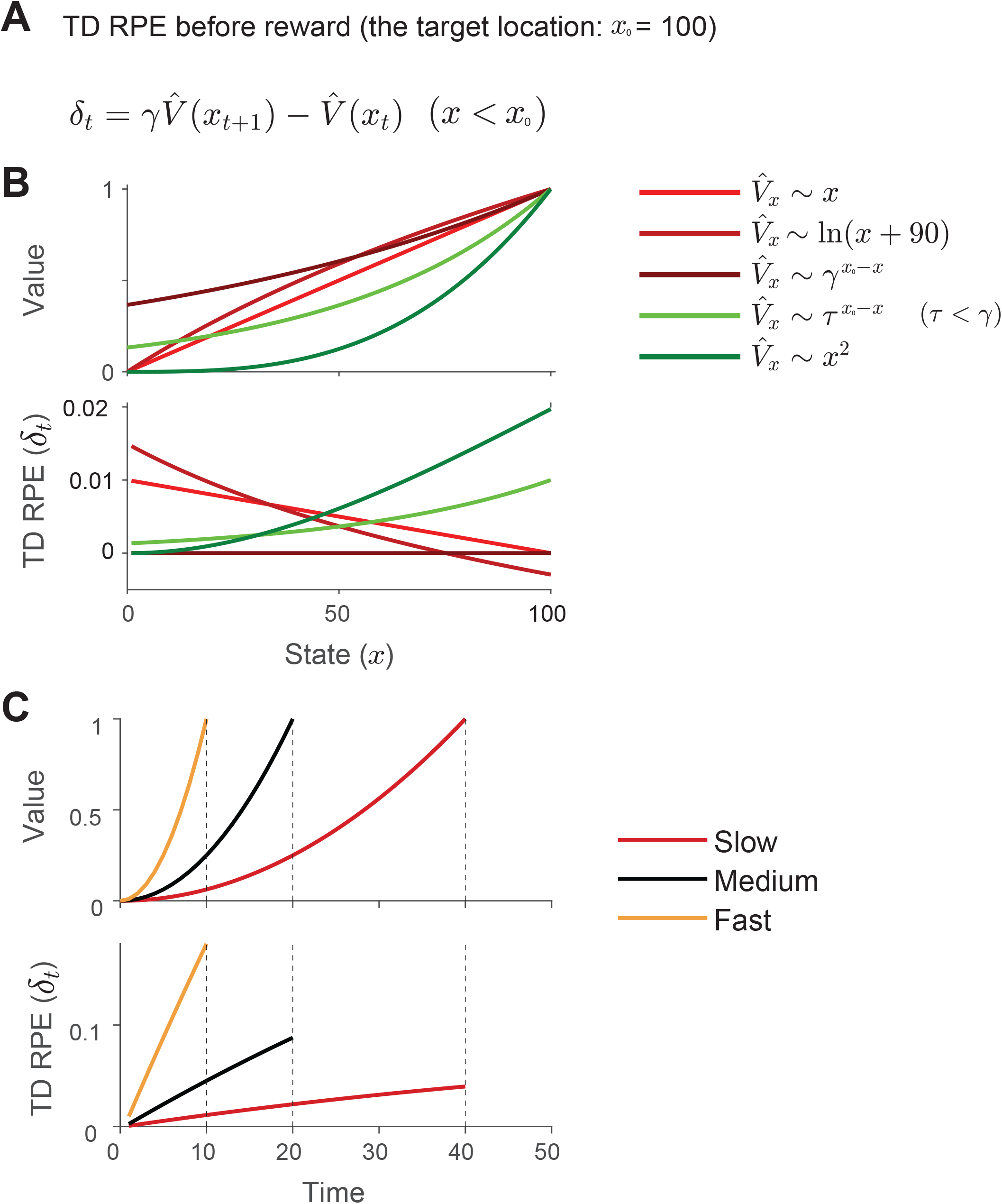
Relationship between value function and TD RPE signal. (**A**) Equation for TD RPE (*δ*) before the reward location 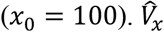, estimated value of state *x*. *γ*, discounting factor (*0 < γ ≤ 1*). (A) Conditions for which TD RPE can ramp up or down. (top) Value functions. (bottom) TD RPEs. Five different forms of value functions and resulting TD RPEs following the equation in (A) are shown. The value functions drawn in green satisfy the ramping condition over the evaluated domain, whereas those drawn in red do not. Note that when the value is discounted by the discounting factor *γ*, TD RPE is zero (dark brown). By contrast, when the value becomes *convex enough* due to a separate spatial discounting factor *τ (τ < γ)*, TD TPE can ramp up (light green). Here we set *γ =* 0.99, and *τ =* 0.98. See Supplementary Note. (**C**) Relationship between speed and TD RPEs. When states are traversed faster, the convexity of the value function will be accentuated when plotted against time. Hence, TD RPEs show greater ramping in faster conditions. Dashed black lines denote the end of the trial for the fast, medium, and slow conditions. Here, we set *γ =* 0.99, track length to 20, and the speeds to 2, 1, and 0.5, for the fast (light orange), medium (black), and slow conditions (red), respectively. Value takes a quadratic shape, and the peak value is normalized to one, i.e., 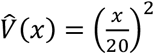

**Fig. S2.**
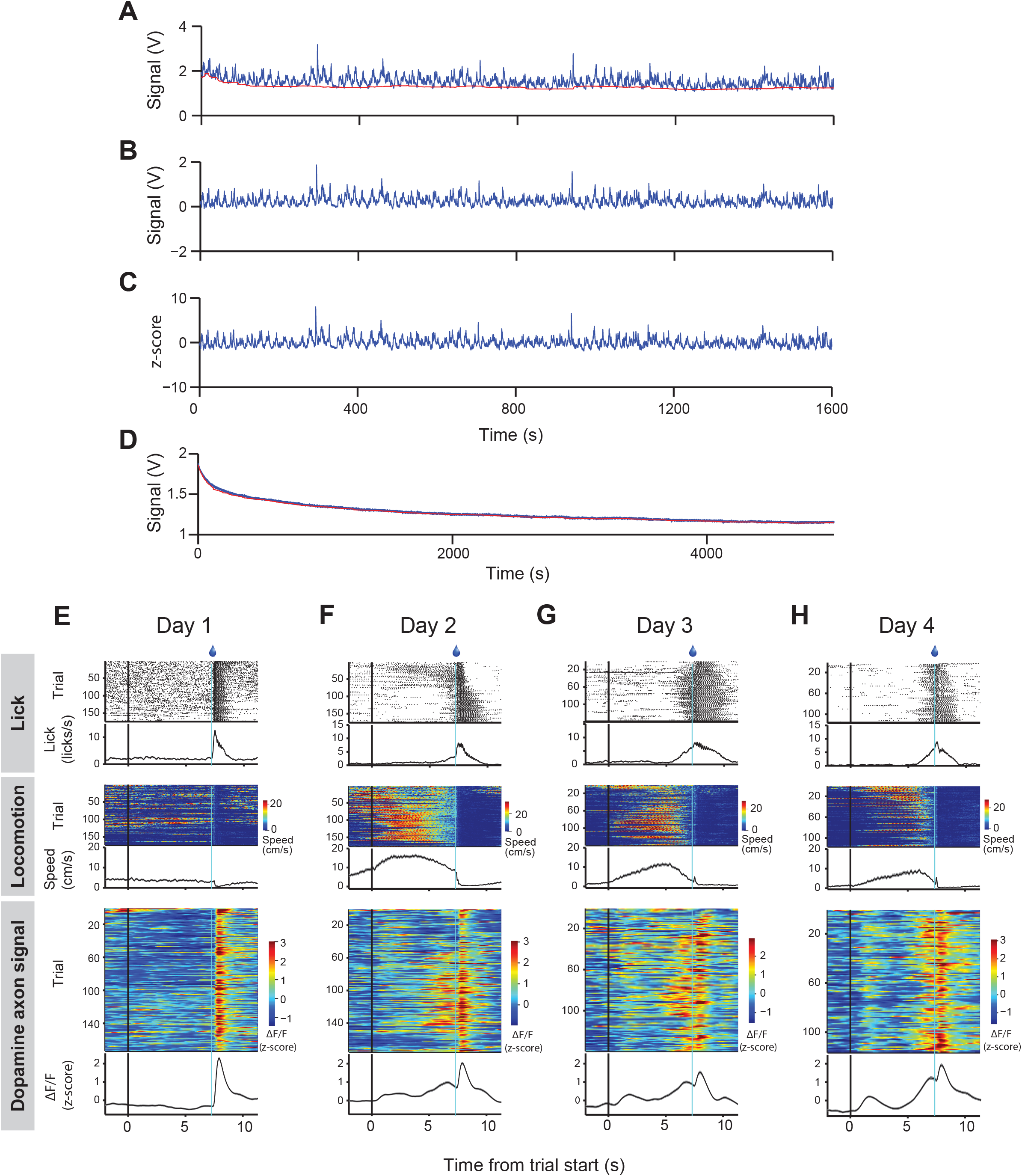
Fluorometry signal processing and example training sessions. (A) Raw voltage outputs from a current amplifier (blue) attached to a photodiode. Slow drift (red) was defined by the lowest 10% of signals using a 2-min moving window. Drift-corrected signal (B) was defined by subtraction of slow drift baseline (A, red) from the raw signals (A, blue). (C) A session-wide mean was subtracted from drift-corrected signals, then the result was divided by the session-wide standard deviation to calculate z-scored fluorescent signals. (D) Raw signals collected from a control animal expressing GFP in dopamine neurons. Signals from the same amplifier gain as (A) are very smooth and lack fluctuations, confirming that motion artifacts were negligible in our head-fixed setup. (**E**) Data collected on day 1 of training. (top) Raster plot showing lick events aligned by scene movement onset. Lick events are averaged across trials using a temporal window of 0.2 s. (middle) Instantaneous speed is color-coded and averaged across trials. (bottom) Z-scored dopamine axon signals are color-coded and averaged across trials. Shading represents s.e.m.. (**F**, **G**, **H**). Data collected on days 2, 3, and 4 (the last day of training for this animal), respectively. Format same as (**E**).

**Fig. S3.**
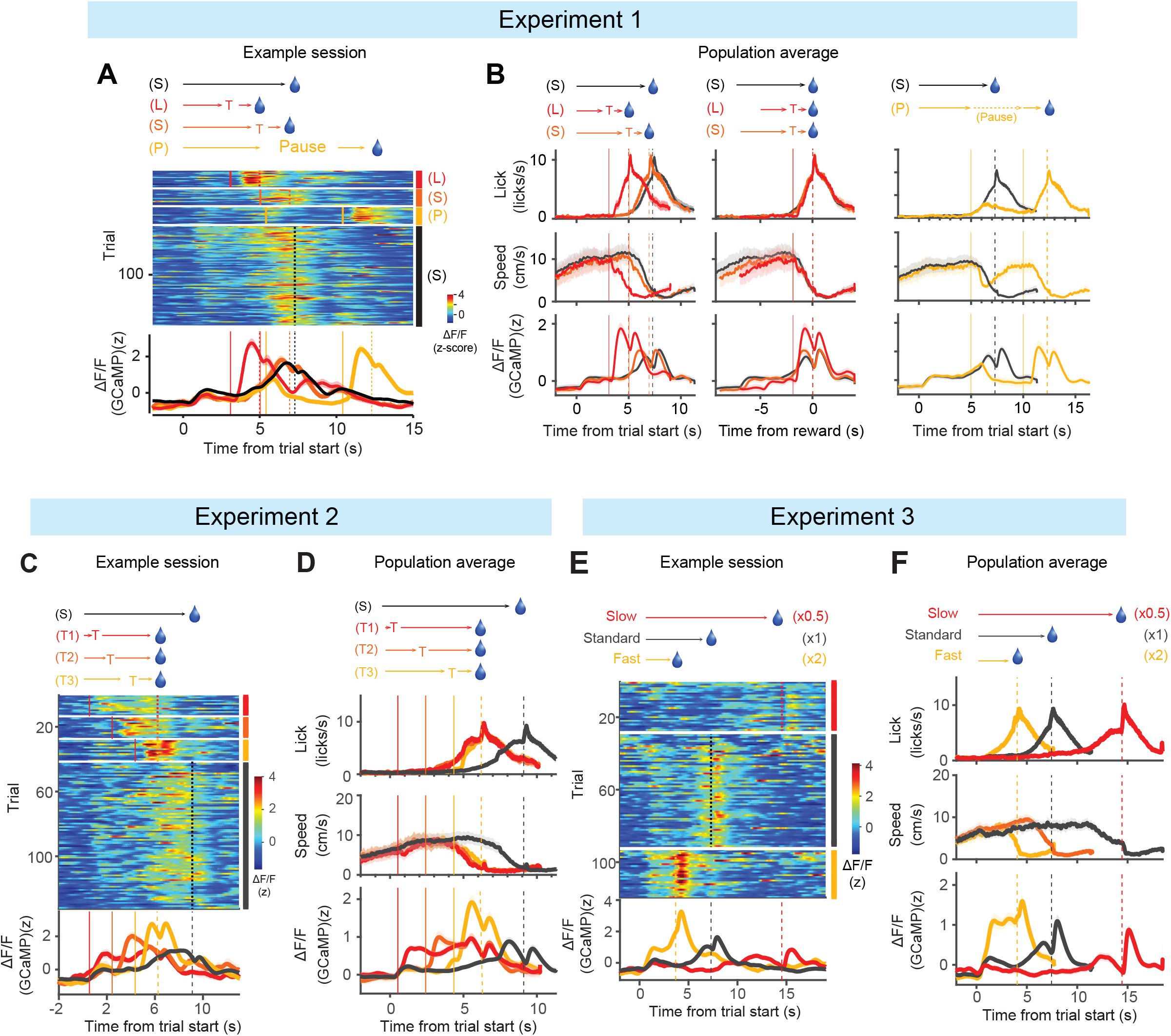
Example calcium recording sessions and population average for Experiments 1, 2, and 3. (**A**) An example session for Experiment 1 (teleport and pause experiment). (top) Time courses of events for each condition (black S: standard; red L: long teleport; orange S: short teleport; yellow P: pause) in Experiment 1. (middle) Z-scored dopamine axon signals from an example session. Trials are sorted by conditions; long-distance teleport (L, red), short-distance teleport (S, orange), pause (P, light orange), and the standard condition (S, black). (bottom) Responses are averaged across trials for each condition. Shading represents s.e.m.. (**B**) Time courses of averaged lick (top row), locomotion speed (middle row), and dopamine axon signals (bottom row) across animals (*n* = 11 mice). Teleport responses are aligned by scene movement onset (left column) or reward onset (middle column). Shading represents s.e.m.. Note that anticipatory lick and slowdown of locomotion overlap when responses are aligned by reward onset, suggesting that animals’ appetitive behaviors were based on their position in the virtual space but not based on the elapsed time alone. (right column) Responses in the pause condition aligned by scene movement onset. (**C**) An example session for Experiment 2 (Three-teleport experiment). Trials are sorted by teleport at short (red), middle (orange), or long (light orange) distance from the start location. Black indicates the standard condition. (**D**) Averaged lick, locomotion speed, and dopamine axon signals (*n* = 11 mice). Teleport onsets (solid line) and water delivery (dashed line) are marked. Shading represents s.e.m..(**E**) an example session for Experiment 3 (Speed-manipulation experiment). Trials were sorted by the speed of scene movement (×0.5: red, ×1: black, ×2: light orange). (**F**) Population-average of lick, locomotion speed, and dopamine axon signals (*n* = 15 mice). Shading represents s.e.m..

**Fig. S4.**
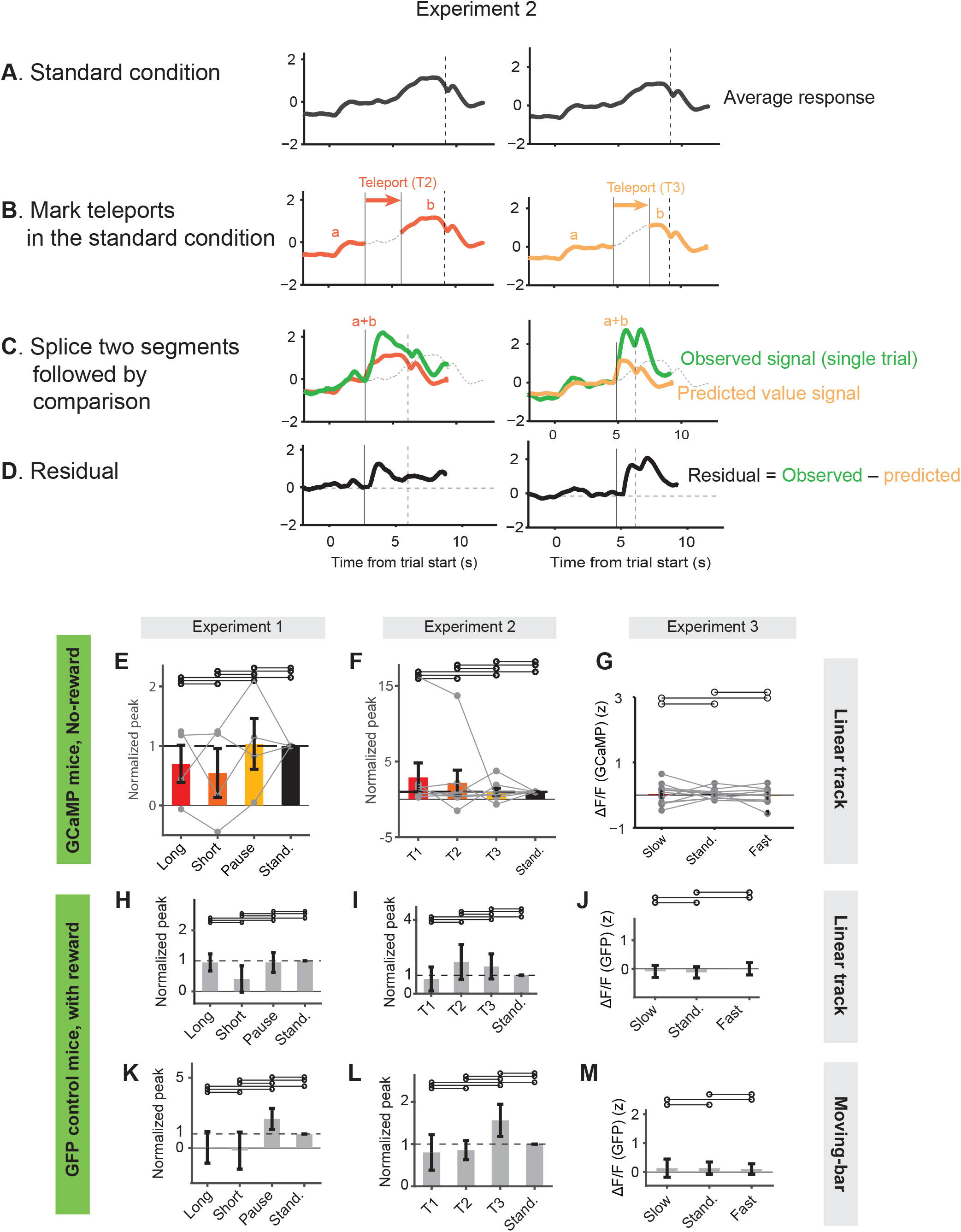
Quantification of residual responses and data from the control conditions. (A-D) Step-by-step procedures for computing residual responses of value model using examples from Experiment 2 (T2 and T3 trials). (**A**) Averaged response in the standard condition (black) was used to predict value signals in the experimental conditions, assuming that the value is a function of location (*x*). (**B**) The start and end locations of a teleport were converted into the time points in the standard condition (black line). Predicted value signals were obtained by splicing (**C**) the segments before and after the teleport timing (a and b, respectively in B). The predicted value signal was then compared with the observed signal. (**D**) The residual was obtained as the difference between the observed and predicted value signals. (**E-G**) Results from no-reward control sessions. Prior to the beginning of training for the standard task, a subset of animals performed Experiments 1-3 as shown in Fig. 2 but without reward at the target location (see Methods). (**E**) Peaks for each condition (*n* = 4, *P* = 1.00, Kruskal-Wallis test). Error bars represent s.e.m.. (F) Normalized peaks for each condition (*n* = 8, *P* = 0.33, Kruskal-Wallis test) in Experiment 2 (three-teleport experiment, corresponding to Fig. 3G). (**G**) Mean responses at [–1 s 0 s] relative to reward onset (*n* = 8, *P* = 0.65, Kruskal-Wallis test) in Experiment 3 (speed-manipulation experiment). (**H-M**) Results from GFP-control animals. Motion artifact in the fluorometry signals was assessed by GFP control animals (See Methods). (**H-J**) Summary of results in Experiments 1-3 in the linear track tasks (*n* = 5 mice). (**K-M**) Summary of results in Experiments 1-3 in the moving-bar tasks (*n* = 4 mice). Error bars represent s.e.m..

**Fig. S5.**
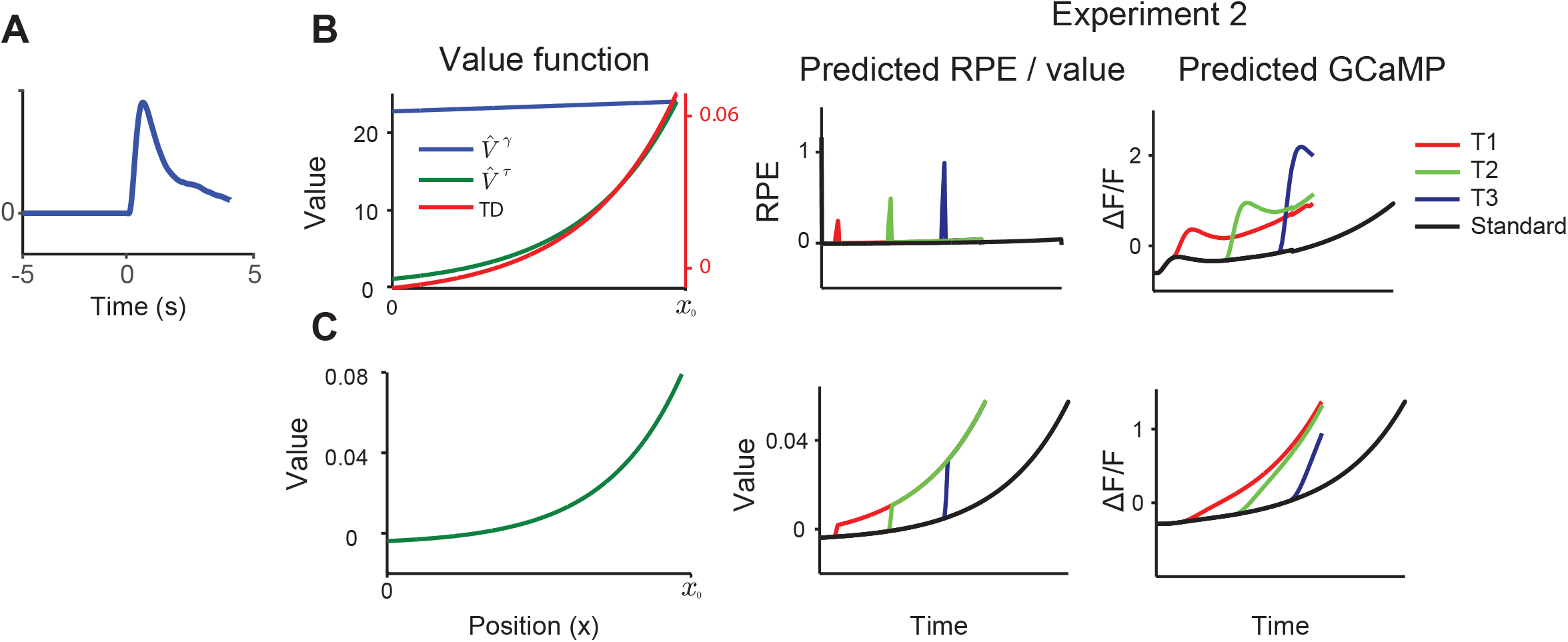
Model fitting analysis. (**A**) GCaMP filter used for convolution. We averaged z-scored responses to unexpected reward in the last training sessions of the standard linear track task (*n* = 17 mice). (**B,C**) Signals predicted by the best fit model for a session of Experiment 2. For this example, the spatial value function was defined by an exponential discounting function 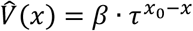. (**B**) (left) The shape of the spatial value function defined by the best fit parameters *τ* and *β* in the RPE model 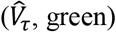. The corresponding TD RPE signals defined by best fit parameters *τ*, *β*, and *γ* in the standard condition (TD, red) show similar shape to the value function. To visualize how convex the shape of the value function is in relation to the temporal discounting factor, the discounting function based on *γ* in the TD equation is also shown 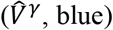 (middle) TD RPE (top) signals during experimental conditions (Experiment 2, T1, T2, T3). Red, T1; green, T2; blue, T3; black, standard. (right) the TD RPE and value signals are then convolved by the GCaMP filter (A) to predict GCaMP (Ca^2+^) signals. (**C**) (left) Value function defined as best fit parameters *τ* and *β* in the value model. (middle) Time courses of value signals. (right) Time courses of predicted calcium signals. Format as top panels.

**Fig. S6.**
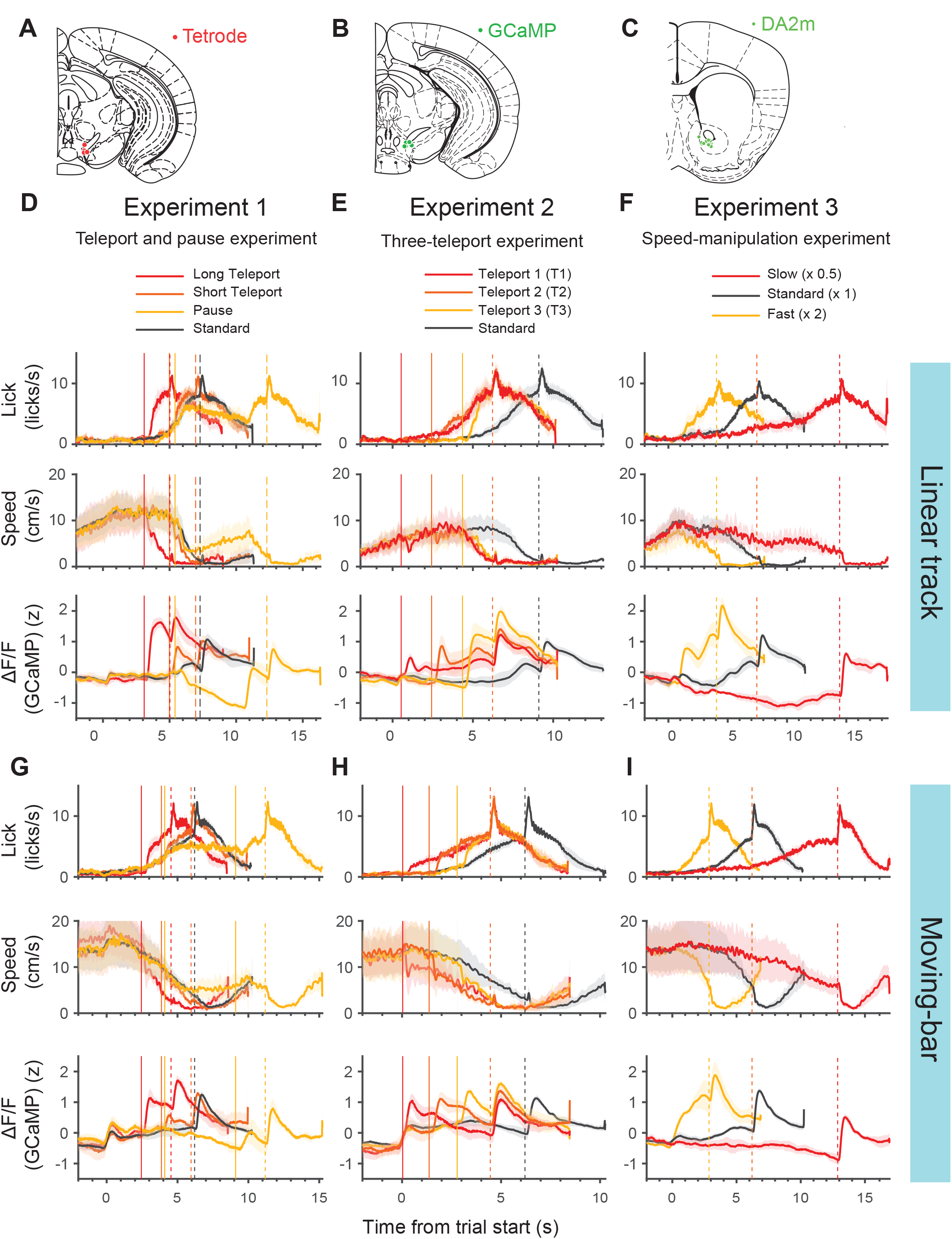
Dopamine cell body activity in the VTA encodes RPE. (**A**) Tetrode positions for single-unit recordings in the VTA (Fig. 5). (**B**) Fiber tip positions for cell body calcium recordings in the VTA. (**C**) fiber tip positions for dopamine concentration in the ventral striatum. (**D-F**) Averaged lick (top), locomotion speed (middle), and fluorometry (photometry) signals in VTA (bottom) in Experiment 1 (**D**), Experiment 2 (**E**), and Experiment 3 (**F**) in the virtual linear track experiments (*n* = 4 mice). Format same as Fig. 3C, G, K, respectively. (**G-I**) Results using moving-bar tasks (*n* = 3). Format same as Fig. 7B, C, D, respectively.

**Fig. S7.**
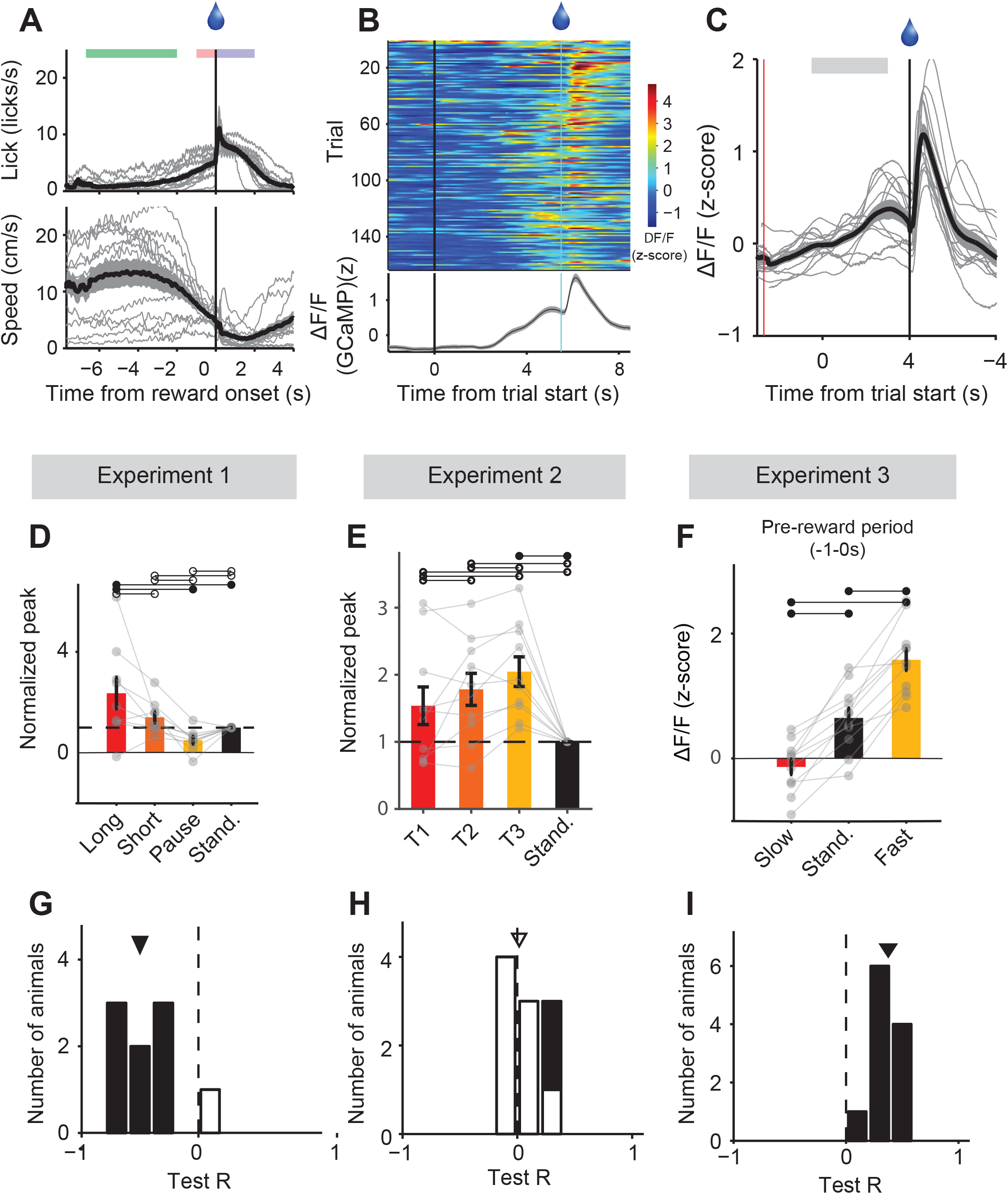
Behaviors and dopamine axon signals in the moving-bar task. (**A**) Time courses of lick rates (top) and locomotion speed (bottom) from individual animals (gray) as well as average across animals (black, *n* = 12 mice). (**B**) Dopamine axon signals from an example session. Shading represents s.e.m.. (**C**) Z-scored dopamine axon signals from individual animals (gray, *n* = 12 mice) and the averaged response (black). Gray horizontal bar represents a time window used to compute Ramping *R*. (**D-I**) Statistical analysis in moving-bar tasks. (**D**) Normalized peak responses from Experiment 1 (Fig. 7B). The median of peaks in the long-distance teleport is significantly greater than 1 (*n* = 9, *P* = 0.039, signed rank test). Error bars represent s.e.m.. (B) Residuals of responses from the state value model prediction are different across conditions (*n* = 9 mice, *P* < 10^-20^, Kruskal-Wallis test). (**E**) Normalized peaks from Experiment 2 (Fig. 7C). (**F**) Results from Experiment 3 (Fig. 7D). Averaged dopamine axon signals from bar movement onset to reward onset. Responses are significantly different (*n* = 11 mice, *P* < 10^-4^, Kruskal-Wallis test). (**G-I**) Summaries of Test *R*s in Experiments 1-3. (**G**) The median *Test R* (black triangle) is significantly less than zero (median *R* = −0.50, *P* = 0.0078, signed rank test). (**H**) The median *Test R* (open triangle) is not different from zero (*P* = 0.43, signed rank test). (**I**) All *Test R*s are significantly greater than zero (alpha = 0.05), and median *R* is significantly greater than zero (median *R* = 0.38, *P* = 0.0010, signed rank test).

## METHODS

### Animals

Twenty-two adult male mice were used for the experiments using calcium indicator (GCaMP). Eighteen mice were heterozygous for the gene expressing the Cre recombinase under the control of the promoter of the dopamine transporter (DAT) gene (Bäckman et al., 2006) (DAT-Cre or B6.SJL-Slc6a3tm1.1(cre)Bkmn/J mice; The Jackson Laboratory; RRID:IMSR_JAX:006660). Four mice were the result of a cross between DAT-Cre mice and a tdTomato transgenic line such that they were heterozygous for DAT-Cre and also heterozygous for tdTomato (Gt(ROSA)26Sortm9(CAGtdTomato)Hze, Jackson Laboratory) (Madisen et al., 2010). We did not observe a difference in results between these mice, thus results were combined. Ten adult C57/BL6J wild-type male mice were used for dopamine sensor experiments. Four adult C57/BL6J DAT-Cre male mice were used for extracellular recording experiments. All mice were backcrossed for >5 generations with C57/BL6J mice. Animals were singly housed on a 12 hr dark/12 hr light cycle (dark from 07:00 to 19:00). All procedures were performed in accordance with the National Institutes of Health Guide for the Care and Use of Laboratory Animals and approved by the Harvard Animal Care and Use Committee.

### Surgery and virus injections

#### Surgery for fiber fluorometry of GCaMP signals

To prepare animals for recording, we performed a single surgery with three key components: (1) AAV-FLEX-GCaMP virus injection into the midbrain, (2) head-plate installation, and (3) implantation of one or more optic fibers into the striatum (Babayan et al., 2018; Menegas et al., 2017). At the time of surgery, all mice were 2–3 months old. All surgeries were performed under aseptic conditions with animals anesthetized with isoflurane (1-2% at 0.5 −1.0 L/min). Analgesia (ketoprofen for post-surgery treatment, 5 mg/kg, I.P.; buprenorphine for pre-operative treatment, 0.1 mg/kg, I.P.) was administered for 3 days following each surgery. We removed the skin above the surface of the brain and dried the skull using air. To express GCaMP specifically in dopamine neurons, we unilaterally injected 250 nl of AAV5-CAG-FLEX-GCaMP6m (1×10^12^ particles/ml, Penn Vector Core) into both the VTA and SNc (500 nl total). To target the VTA, we made a small craniotomy and injected virus at bregma 3.1, lateral 0.6, depths 4.4 and 4.1 mm. To target SNc, we injected virus at bregma 3.3, lateral 1.6, depths 3.8 and 3.6 mm. Virus injection lasted several minutes, and then the injection pipette was slowly removed over the course of several minutes.

We then installed a head-plate for head-fixation by gluing a head-plate onto the top of the skull (C&B Metabond, Parkell). We used ring-shaped head-plates to ensure that the skull above the striatum would be accessible for fiber implants. Finally, during the same surgery, we also implanted optic fibers into the ventral striatum. For a subset of animals, we also implanted fibers into dorsomedial or dorsolateral striatum. To do this, we first slowly lowered optical fibers (200 µm diameter, Doric Lenses) into the striatum using a fiber holder (SCH_1.25, Doric Lenses). The coordinates we used for targeting were bregma 1.0, lateral 1.1, depth 4.1 mm. Once fibers were lowered, we first attached them to the skull with UV-curing epoxy (Thorlabs, NOA81), and then a layer of black Ortho-Jet dental adhesive (Lang Dental, IL). After waiting for fifteen minutes for this glue to dry, we applied a small amount of rapid-curing epoxy (A00254, Devcon) to attach the fiber cannulas to the underlying glue and head-plate. After waiting for fifteen minutes for the epoxy to cure, the surgery was completed.

#### Surgery for fiber fluorometry of dopamine sensor signals

Surgical procedures up to virus injection were the same as GCaMP injections described above. Instead of GCaMP, we injected 400nl of AAV9-hSyn-DA2m (Vigene bioscience) into the ventral striatum (bregma 1.0, lateral 1.1, depths 4.2 and 4.1mm).

Once injection was completed, we implanted fiber and head plate in the same way as the surgery for GCaMP recording.

#### Surgery for single unit recording of opto-tagged dopamine neurons

We performed two surgeries, both stereotactically targeting the left VTA (from bregma: 3.1 mm posterior, 0.6 mm lateral, 4.2 mm ventral). In the first surgery, we injected 500 nl of adeno-associated virus serotype 5 (AAV5) carrying an inverted ChR2-encoding sequence (H134R) fused to the sequence expressing the fluorescent reporter eYFP and flanked by double *loxP* sites3,37. We previously showed that the expression of this virus is highly selective and efficient in dopamine neurons3. After 2 weeks, we performed the second surgery to implant a head plate and custom-built microdrive containing 8 tetrodes and an optical fiber.

### Virtual reality setup

Virtual environments were displayed on three liquid crystal display (LCD) monitors with thin frames (width 53 cm, height 30 cm) that were placed on the left, front, and right side of animals (Chen et al., 2013a; Harvey et al., 2009). A workstation computer (DELL Precision 5810) with a high-performance graphics card (NVIDIA Quadro K2200) was used to present visual images. VirMEn software (Aronov and Tank, 2014) was used to generate virtual objects and render visual images using perspective projection. No dropping of image frames was confirmed by measuring intervals in the inter-frame callback function, and by photodiode measurements while a test program alternated the brightness of the screen every frame.

Animals were head-restrained at the center of three monitors, 7.5 cm above the bottom of the screen. Mice were placed on a cylindrical styrofoam treadmill (diameter 20.3 cm, width 10.4 cm) (Howe and Dombeck, 2016). The rotational velocity of the treadmill was encoded using a rotary encoder. The output pulses of the encoder were converted into continuous voltage signals using a customized Arduino program running on a microprocessor (Teensy 3.2).

Water reward was given through a water spout located in front of the animal’s mouth. Licking tongue movements were monitored using an infrared sensor (OPB819Z, TT Electronics). Voltage signals from the rotary encoder and the lick sensor were digitized into a PCI-based data-acquisition system (PCIe-6323, National Instruments) installed on the visual stimulation computer. Timing and amount of water were controlled through a micro-solenoid valve (LHDA 1221111H, The Lee Company) and switch (2N7000, On Semiconductor). Analog output TTL pulse was generated from the visual stimulation computer to deliver reward to the animals.

### Virtual linear track experiments

Animals were trained in a virtual linear maze (Fig. 1, length of 100 a.u. and width of 30 a.u.). The maze was composed of a starting platform and a corridor with walls on both sides. The walls have four different texture patterns to help animals recognize a position in the virtual space (Supplementary Movie 1).

We first trained animals on the standard approach-to-target task to learn the association between target location and reward. Once the animals learned the task, we ran a series of tasks with ‘probe’ trials to examine the nature of dopamine signals. We typically ran each task (Experiments 1-4) for two consecutive days (with zero-or one-day break). A daily session started with 10 standard trials to help animals remember the task before presenting any probe trials. Unless otherwise noted, unexpected reward was given during the inter-trial interval (5 µl) on 3-6% of trials.

#### Standard approach-to-target task

The session started in a dark gray background. Trials started with the presentation of the visual scene with the animal placed at the starting position (0 arbitrary units, a.u.). After a random delay (1-s offset plus a random delay drawn from a modified exponential distribution with a mean of 1.5 s and cutoff of 3.5 s), the visual scene started moving forward. The velocity linearly increased for 1 s until it reached 13 a.u./s, after which the velocity was maintained to be constant until the animal reached the target position (97 a.u.). A drop of water reward (5 µl) was delivered through a water spout that was placed in front of the animal’s mouth. No reward omission trials were used. Once the reward was given, the visual stimulus was turned off after a random delay (drawn from an exponential distribution with a mean of 1 s, to which a 1-s offset was added). Delay was re-drawn if it exceeded 4 s. Inter-trial interval (ITI) was drawn from an exponential distribution with a mean of 3 s, to which a 3.5-s offset was added. ITI was re-drawn if it exceeded 10 s.

#### Experiment 1 (Teleport to the same destination Task)

The following three types of probe conditions were randomly interleaved with the standard condition. Each probe condition comprised 20% of trials.

1. *Long teleport*: when the animal reached a predefined position (40 a.u.), it was teleported to a position closer to the reward (70 a.u.). At the time of teleport, the screens were briefly (93 ms) blanked to black.
2. *Short teleport*: when the animal reached another predefined position (65 a.u.), it was teleported to the same destination (70 a.u.).
3. *Pause*: when the animal reached this destination (70 a.u.), the progression of the scene was paused for 5 s, after which the scene movement resumed.

In all trials, the animal received a reward when it reached the same goal location (97 a.u.).

#### Experiment 2 (Three-teleport Task)

The following three types of teleport conditions were randomly interleaved with the standard condition with the frequency of all teleport conditions being 33-40% in total. On a fraction of trials, when the animal reached one of the three teleport positions (5, 25, or 45 a.u.), it was teleported by the same distance (30 a.u.) forward to the location closer to the goal location (35, 55 and 75 a.u., respectively). The screens were briefly (93 ms) blanked to black at the time of teleport. Scene progression was 20% slower than that it had been during training, except for the first four animals. Data from these four animals were excluded from the population PSTH but included in other statistical analyses.

#### Experiment 3 (Speed-manipulation Task)

On a fraction of trials (20%), the progression of the scene was twice as fast as in the standard condition. On some other trials (20%), the progression was half as fast as in the standard condition. The rest of the trials were the same as the standard approach-to-target task. The acceleration of the scene in the manipulated condition was identical to the standard condition (13 a.u./s^2^).

#### Experiment 4 (Reward-size Varying Task)

We alternated blocks of 25 trials to switch between a small-reward (2.25 µl/trial) and a big-reward (10 µl/trial) context. No explicit cue was given to notify the block switch. On the first day in Experiment 4, we randomly chose whether the first block was a small- or big-reward block. The other reward size was used for the first block on the next day. On a fraction of trials (20%), we teleported the animals from 45 a.u. to 75 a.u..

#### No Reward control experiments

We tested whether the transient response to teleport is due to the short blackout or the abrupt change of visual scene in the absence of a reward context. Before training began with reward in the standard approach-to-target condition, a subset of animals performed the same-destination (*n* = 7 mice), three-teleport (*n* = 13 mice), and speed-manipulation (*n* = 13 mice) conditions without giving reward at the target location. Unexpected rewards were delivered during the inter-trial intervals on 10% of trials. We ran one session for each protocol (70-100 trials), one or two control experiments a day.

### Moving-bar experiment

We used the same three-monitor display setup as in our virtual reality task. The background was gray. A black ring-shaped object was used to render black bars on the three screens in the virtual environment. The object (2.5 cm vertical thickness) moved vertically at a constant speed to indicate reward proximity.

#### Standard task

A black bar was initially presented at the top of the screen. After a random inter-trial interval, the bar started to move from the top to the bottom of the screen at a constant speed (3.7 cm/s). When the bar reached a goal position (25 cm from the top of the screen, 6.7 s after the movement onset of the bar), a drop of water (5 µl) was delivered. After a random delay (the same as the 1D maze task), the bar was shifted back to the original starting position. The screen was kept on during the inter-trial interval.

#### Experiment 1: Two teleport and one pause task

On some fraction of trials (12.5%, respectively), when the position of the bar reached one of two positions on the screen (10.9 cm or 16.25 cm from the top of the screen), the bar was abruptly shifted downward by 6.25 cm or 0.93 cm, respectively, and maintained movement with constant speed. On another 12.5% of trials, the bar movement was paused for 5.0 s, after which the movement resumed. These probe conditions were randomly interleaved with the standard condition.

#### Experiment 2: Three-teleport task

When the center of the bar reached 1.56, 6.9, or 12.18 cm from the top of the screen, the bar shifted its position by 6.6 cm (12.5% of total trials, respectively). The teleport conditions were randomly interleaved with the standard condition.

#### Experiment 3: Speed manipulation task

The movement of the bar was twice as fast as the standard condition (20%) or half as fast as the standard condition (20%).

### Fiber fluorometry (photometry)

Fluorescent signals from the brain were recorded using a custom-made fiber fluorometry (photometry) system as described in our previous studies (Babayan et al., 2018; Menegas et al., 2017). The blue light (473 nm) from a diode-pumped solid-state laser (DPSSL; 80–500 μW; Opto Engine LLC, UT, USA) was attenuated through a neutral density filter (4.0 optical density, Thorlabs, NJ, USA) and coupled into an optical fiber patchcord (400 μm, Doric Lenses) using a 0.65 NA microscope objective (Olympus). The patchcord connected to the implanted fiber was used to deliver excitation light to the brain and to collect the fluorescence emission signals from the brain. The fluorescent signal from the brain was spectrally separated from the excitation light using a dichroic mirror (T556lpxr, Chroma), passed through a bandpass filter (ET500/50, Chroma), focused onto a photodetector (FDS100, Thorlabs) and amplified using a current preamplifier (SR570, Stanford Research Systems). Acquisition from the red fluorophore (tdTomato) was simultaneously acquired (bandpass filter ET605/70 nm, Chroma) in some animals (*n* = 4 mice) but was not used for further analyses. The voltage signal from the preamplifier was digitized through a data acquisition board (PCI-e6321, National Instruments) and stored in a computer using a custom software written in LabVIEW (National Instruments).

Motion artifacts were examined using mice expressing GFP in dopamine neurons. We injected AAV5-FLEX-GFP in the VTA and SNc of DAT-Cre mice and collected behavior and fluorometry signals in the same way as the GCaMP animals. Raw voltage traces were qualitatively different from GCaMP, with no noticeable fluctuation with the same amplifier configuration. Although animals learned the tasks and acquired behaviors similar to the GCaMP animals, we observed no ramping. We observed neither phasic responses to teleports, nor any significant difference between test conditions.

### Electrophysiology

We based recording techniques on previous studies (Cohen et al., 2012; Kvitsiani et al., 2013; Lima et al., 2009; Tian and Uchida, 2015). We recorded extracellularly from the VTA using a custom-built, screw-driven containing eight tetrodes glued to a 200-µm optic fiber (ThorLabs). Tetrodes (Sandvik, Palm Coast, Florida) were glued to the fiber and clipped so that their tips extended 200–500 µm from the end of the fiber. We recorded neural signals with an Open Ephys recording system with Intan headstage (RHD2132, Intan technologies). Broadband signals from each wire were recorded continuously at 30 kHz. To extract spike timing, signals were band-pass-filtered between 300 and 6,000 Hz and sorted offline using MClust-4.3 (A.D. Redish). To be included in the data set, a neuron had to be well isolated (a measure of unit isolation quality, L-ratio < 0.05) (Schmitzer-Torbert and Redish, 2004). We also histologically verified recording sites by creating electrolytic lesions using 10–15 s of 30 µA direct current.

To unambiguously identify dopamine neurons, we used ChR2 to observe laser-triggered spikes (Cohen et al., 2012; Lima et al., 2009). The optical fiber was coupled with a diode-pumped solid-state laser with analog amplitude modulation (Laserglow Technologies). At the beginning and end of each recording session, we delivered trains of ten 473 nm light pulses, each 5 ms long, at 1, 5, 10, 20 and 50 Hz, with an intensity of 5–20 mW/mm^2^ at the tip of the fiber. To be included in our data set, neurons had to fulfill three criteria. (i) The neurons’ spike timing must be significantly modulated by light pulses. We tested this by using the stimulus associated spike latency test (SALT) (Kvitsiani et al., 2013). We used a significance value of *P* < 0.05, and a time window of 10 ms after laser onset. (ii) Laser-evoked spikes must be near-identical to spontaneous spikes. This ensured that light-evoked spikes reflected actual spikes instead of photochemical artifacts. Neurons must have a short latency to spike following laser pulses, and little jitter in spike latency.

### Histology

Mice were perfused with phosphate buffered saline (PBS) followed by 4% paraformaldehyde in PBS. The brains were cut in 100-µm coronal sections using a vibratome (Leica). Brain sections were loaded on glass slides and stained with 4′,6-diamidino-2-phenylindole (DAPI, Vectashield). The locations of fiber tips were determined using the standard mouse brain atlas (Franklin and Paxinos, 2008).

## DATA ANALYSIS

### Fluorometry (photometry)

Power line noise in the raw voltage signals was removed by notch filter (MATLAB, Natick, MA). A baseline of the voltage signal was defined by the lowest 10% of signals using a 2-min window. The baseline was subtracted from the raw signal, and the results were z-scored by a session-wide mean and standard deviation (Fig. S2). For PSTHs in Experiment 4 using fiber fluorometry, the fluorescent level from 1-0 s before trial onset was subtracted. Both GCaMP signals and dopamine sensor signals were processed in the same way.

### Licking and locomotion

Animals lick the water spout in anticipation of water delivery. To measure licking, we used a photoelectric sensor that output a voltage based on the disruption of its infrared light path (OPB819Z, TT Electronics). Lick timing was defined as deflection points (peaks) of the output signals above a threshold. To plot the time course of licks, instantaneous lick rate was computed by a moving average using a 200-ms window.

We used the following time window to quantify the lick rate. Impulsive lick: from visual scene movement onset to 2 s before reward; anticipatory lick: [-1 s 0 s] relative to reward; post-reward lick: [0 s 2 s] relative to reward. The same temporal windows are used for quantifying speed. Net anticipatory licking and anticipatory slowdown are defined as below:

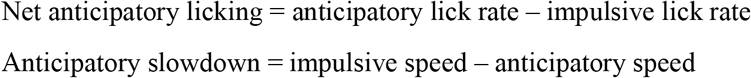

### Dopamine ramping (Ramping R)

We used the trials in the standard condition to quantify the ramping of fluorescent signals. We computed the Pearson correlation coefficient between time points and z-scored fluorescent signals using either individual trials or averaged response. For the linear track tasks, we used the time window of [-3.5 s −1 s] relative to reward onset for fiber fluorometry data and [-5 s −1 s] for spiking data. We used a smaller window for fluorometry as the slowly delaying response to stimulus or movement onsets often biases the slope of the ramping. For the moving-bar tasks, we used the time window of [-4.5 s −1 s] relative to reward onset.

### Session-averaged time course

Lick, locomotion speed, and z-scored dopamine responses for individual trials were aligned by external events (e.g., trial start or teleport onset), and then smoothed using a moving average method. We used a 200ms time window for licking and spike signals. We did not smooth locomotion speed and fluorometry signals. The results were then averaged across trials for each experimental condition to generate a session-averaged time course.

### Population-averaged time course

For calcium recording experiments, we computed the mean of session-averaged time courses from the second session dataset (as the average of all session averages) along with the standard error (the total number of sessions being the sample size) for each experimental condition. Population-average time courses are used to summarize behavior and dopamine responses. Since we did not observe a significant difference in Ramping *R* or Test *R* between the first session and the second session in the calcium data (data not shown), we used both sessions for the dopamine sensor data in plotting population PSTHs.

### Summarizing responses using normalized peak

Responses to teleport and pause were quantified by peaks of session-averaged time courses using time windows [0.6 s – 2.1 s] and [2 s – 5 s] relative to the teleport and pause onset, respectively. Peaks in the test conditions were normalized by peaks in the standard condition using time window from trial start to reward onset (Fig. 3D left, **3H** left).

### Residual responses based on the state value model

In the teleport conditions, forward scene movement was maintained before and after teleports. As a result, phasic responses to teleports can be contaminated by ramping. For more accurate model comparisons, we generated model responses based on state value predictions (Fig. S4A-D). The process begins with a session-averaged time course in the standard condition. Model responses were shifted in time by the amount shortened by teleport (2.2 s for long teleport, 0.25 s for short teleport) or lengthened by pause (5 s). The results were further delayed by 0.3 s to account for neural latency in GCaMP signal. The residual responses were then defined as the subtraction of model response from empirical responses. Deviation from value model was defined as average residual response using time windows [0.6 s – 2.1 s] and [2 s – 5 s] relative to the teleport and pause onset, respectively (Fig. 3D right, **3H** right). The baseline activity for each trial, defined by average response using time window [-1 s – 0 s] from trial start, was subtracted from the average residual responses.

### Summarizing responses for single unit data

For individual trials, responses to reward, teleport, pause were quantified by averaging responses using time windows [0.05 s – 0.45 s], [0.1 s – 0.5 s], [0.1 s – 0.5 s] relative to the events of interest, respectively. Baseline responses were defined by average responses using time window [-1 s – 0 s] relative to trial start. The baseline responses were subtracted from the responses of interests to obtain a net modulation by the events.

### Estimation of GCaMP signals from spiking data

We estimated GCaMP responses of a single neuron based on a relationship between a single spike and GCaMP response (Chen et al., 2013b). We convolved spike train with the GCaMP kernel to plot estimated GCaMP signals from a single neuron activity (Fig. 5H, I, J, bottom). GCaMP signals measured from fiber fluorometry can be approximated by calcium signals pooled across neurons. To estimate the fluorometry response of a single trial, we randomly selected single trials of convolved responses in the same experimental condition and summed up the responses across neurons. We repeated this for multiple trials (*n* = 100 for Fig. 5F; n = 200 for Fig. 5K-M) to generate a predicted GCaMP PSTHs.

### Summarizing responses in the test condition (Test R)

The systematic deviation from value model across conditions was quantified using Spearman correlation for each session. For Experiment 1 (teleport and pause), numbers were assigned to each condition to quantify the trial-by-trial correlation between test conditions and residual responses based on the value model (long teleport = 1; short teleport = 2; pause = 3). Significant negative correlations indicate that residual responses are big in the long teleport, medium in the short teleport, and the smallest in the pause condition. For Experiment 2 (three teleports), we computed Spearman correlations between position and the residual responses. Positive correlations indicate that responses to teleports increase with proximity to reward location. For Experiment 3, we computed Spearman correlation between speed condition and baseline-corrected (using time window [-1 s −0 s] from trial start) average response from trial start to reward onset. Numbers were assigned such that positive correlations indicate that responses increase with scene speed (slow speed = 1; standard speed = 2; fast speed = 3). In addition to testing the significance of trial-by-trial correlation in an individual session, we further tested whether the median of Test Rs across animals is significantly different from zero using signed rank test.

### Statistical analysis

We performed statistical analyses at both single-session and population levels. For individual-session analysis, average responses from individual trials were quantified using a temporal window locked to an external event (e.g., reward onset). Non-parametric tests (e.g., signed rank test) were used to test whether responses are significantly greater or smaller than reference (e.g., zero). A dataset with a significant difference was marked by a filled circle. For population-level analysis, the mean responses of individual sessions were used for comparisons between conditions or comparisons against a reference value (e.g., zero or one) using a signed rank test. A significant difference between conditions was marked by a filled circle on the horizontal lines that connect the two conditions. A significant difference from baseline was marked using a p-value or a star mark on top of each condition. We used two-tailed tests for all statistical tests.

### Across-session regression analysis

We examined whether and how much task-related behaviors can account for the variability in Ramping *R* across sessions. For each session, we quantified average Ramping *R*, net anticipatory lick rate, and running speed. We regressed Ramping *R* on net anticipatory lick rate and running speed across sessions, including the last day of training, Experiments 1, 2, and 3. For Experiments 1-3, only trials in the standard condition were used.

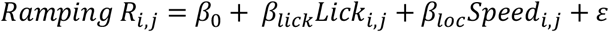

The confidence interval of *β_lick_* and *β_lac_* was used to determine the significance of relationship between individual behavioral variable and Ramping *R*.

### Trial-to-trial regression analysis for Experiment 3

We examined how trial-to-trial variability in the mean dopamine responses during approach can be accounted for by visual scene speed, locomotion speed, and a global trend by multiple linear regression (Fig. 2L, right).

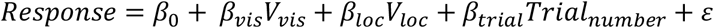

### Model fitting

For quantitative comparisons between the state value and RPE models (Fig. 4), we examined which model provides a better fit to the mean calcium signals from each animal. For this analysis, we focused on the pre-reward period where ramping dopamine signals were observed. We first defined the shape of the value function across space. We then predicted how the value or TD RPE changes in each experimental condition. We then converted the predicted value or TD RPE into calcium signals, and these predictions were compared with the mean calcium signals in the data.

The TD RPE at time *t* (*δ_t_*) is defined by:

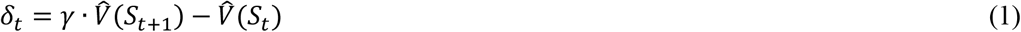

where *S_t_* is the state at time 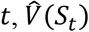 is the estimated value of the state *S_t_*, and *γ* is the discounting factor (*0 < γ ≤ 1*). Note that this formulation excludes reward delivery times. To fit responses in the linear track tasks, we used the position along the linear track as *S_t_*; that is, the state value is defined as the value of the position. To fit responses in the moving-bar tasks, we used the vertical position of the bar as *S_t_*.

The state value is expected to increase as the animal gets closer to the goal location. For the TD signal to ramp up, the value function must take a convex shape along the relevant dimension along which the animal traverses (See Supplementary Note). Because the exact shape of the state value is unknown, we examined several shapes of value functions.

Our first model defines the shape of the value function as an exponentially decaying function across space with the discounting factor *τ*.

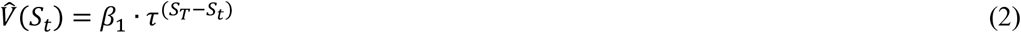

where *β_1_* is a coefficient representing the value at the goal location while *S_T_* is the position of the goal (target), and therefore, *S_T_ − S_t_* corresponds to the distance from the current position to the goal. Note that the discounting factor *τ* in Equation (2) need not be the same as the discounting factor *γ* for the definition of TD errors (i.e., Equation (1)).

Additionally, we explored various other types of functions across space. For instance, we found that a good fit was obtained using a cubic function defined as follows:

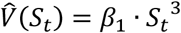

After defining a value function, TD RPE was calculated using Equation (1).

Next, the predicted value or TD RPE was convolved with a kernel (*F_GCaMP_*) to reflect the slow kinetics of calcium signals compared to spikes. We obtained the kernel for GCaMP by averaging GCaMP responses to a delivery of unexpected reward during the last day of training. For the TD RPE model, the calcium signals *y_t_* were predicted by convolving *δ_t_* with the GCaMP kernel and an offset term.

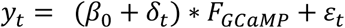

where ε*_t_* denotes Gaussian-distributed random error. For the value model, calcium signals were predicted by convolving 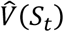 with the GCaMP kernel.

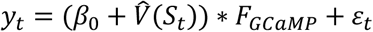

A model fit was performed by minimizing the sum of squared residuals (SSR) in the time window from 2 s after the scene movement onset to 0.5 s before reward delivery. To find parameters that minimize SSR numerically, we used a non-linear function solver with constraints (fmincon, MATLAB). Sets of the example starting point (p0), lower bound (LB) and upper bound (UB) are as follows:

- TD model, exponential 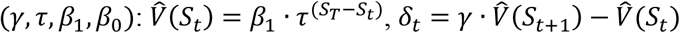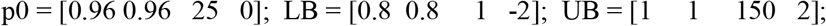
- TD model, cubic 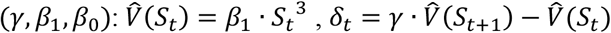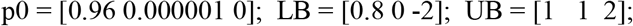
- Value model, exponential 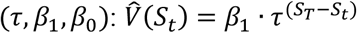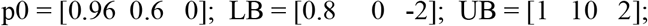

Starting points of parameter search were either the example starting point (p0, n = 1) or randomly drawn from a uniform distribution ranging between the lower and upper bounds (n = 99). The solver was repeated 100 times, and the parameters with the minimum SSR were chosen.

Empirical value functions may deviate from a simple mathematical form (e.g., exponential). To model more general shapes of the value function, we also used third order polynomial regression.

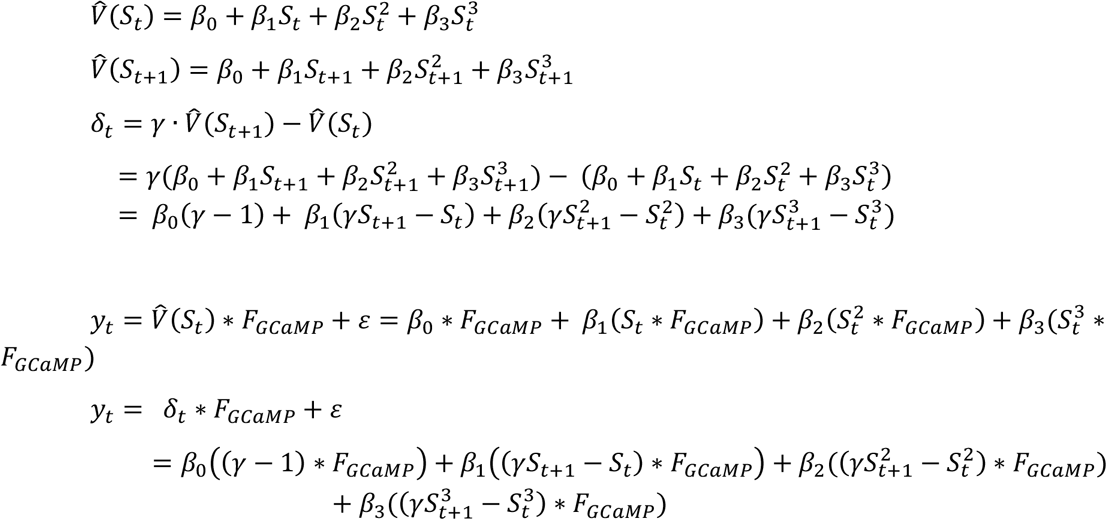

The coefficients of the value model can be found deterministically. For the TD model, we performed regression using a range of *γ* values from 0.97 to 1 in steps of 0.0005 and used the *γ* value with maximum *R^2^*.

To compare goodness-of-fits across models, Akaike information criterion (AIC) (Akaike, 1973) was used.

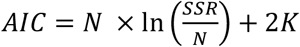

where *N* is the number of data points, SSR is the squared sum of residuals, *K* is the number of fitting parameters. AIC was computed using the time window from 2 s after the scene movement onset to 0.5 s before reward delivery. This window was chosen to exclude phasic responses at the movement scene onset or reward delivery which may bias our model selection to favor RPE models. When computing AIC for polynomial fits, the number of parameters (*K*) was increased by one to compensate for the exhaustive search for *γ*. For comparisons, we computed differential AIC between a target model and a reference model *(ΔAIC = AIC_target_ − AIC_ref_)*. Positive value indicates that the reference model explains the data better than the target model considering the number of parameters.

## SUPPLEMENTARY NOTE

### Temporal difference reward prediction errors are approximately the temporal derivative of the value function, plus received rewards

In reinforcement learning theories (Sutton, 1988), the value of a given state is defined as the sum of all future rewards, where rewards are discounted by a constant rate (*γ*, discounting factor, *0 < γ ≤ 1*) per unit time:

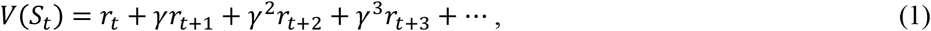

where *r_t_* is the reward at time *t*, *S_t_* is the state at time *t*, and *V(S_t_)* is the value of the state *S_t_*. Under the assumption that state transitions and rewards follow a Markov process, Equation (1) can be rewritten as:

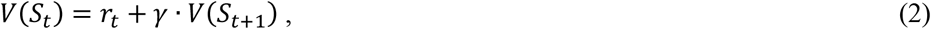

which is known as the Bellman equation (Bellman, 1954). The agent approximates the true value *V(S_t_)* with a learned estimate 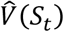, so that if true value is perfectly learned, i.e., 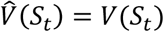, then

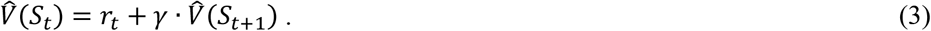

However, before the agent has learned the true value, the left-hand and right-hand sides of Equation (3) will not be equal on average. The difference between these two terms represents the error in value prediction, and as such defines the temporal difference reward prediction error (TD RPE, or *δ*):

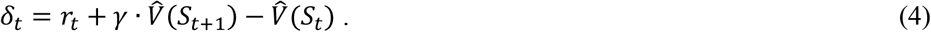

According to this definition, the TD RPE contains a difference between the estimated values of states that are evaluated at consecutive time points, 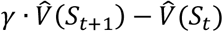. When *γ = 1,* this term is exactly the temporal derivative of the estimated value function. When *γ* is not 1 but close to it, this term is *approximately* the derivative of estimated value. Thus, the TD RPE is approximately the derivative of estimated value, plus rewards received *(r_t_)*(Gershman, 2014). As a result of this property, unexpected increases and decreases in value result in positive and negative transient (“phasic”) changes in the TD RPE, respectively.

TD RPEs, as defined by Equation (4), account for three features of dopamine responses in simple classical conditioning paradigms (Schultz et al., 1997):

1. Dopamine neurons are excited by a cue that predicts future reward. In TD models, this occurs because the reward-predicting cue indicates that value at the time of cue presentation is greater than originally expected (i.e., the animal now expects that a reward is coming).
2. When a predicted reward is omitted, dopamine neurons transiently reduce their firing below baseline. In TD models, this occurs because value at the time of omitted reward is now less than originally expected.
3. Unpredicted rewards excite dopamine neurons. However, when a cue predicts delivery of reward, dopamine neurons’ response to the predicted reward is greatly reduced. In TD models, this occurs because excitation due to received reward is canceled out by the negative response in 2.

### Reward prediction error will ramp when value takes a sufficiently convex shape across time

Consider trials in which a single reward is presented at time *T*. Then before reward is received *(t < T)*, the TD RPE is simply

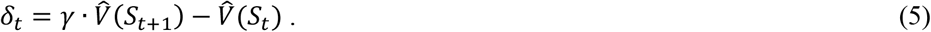

How must value vary with time in order to produce ramping TD RPEs? We can examine this question by writing the necessary and sufficient condition for a monotonic increase in TD RPEs, i.e.,

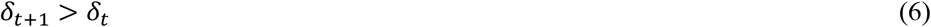

for all *t < T*. Expanding Inequality (6),

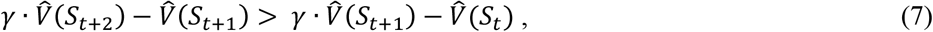

which can be rewritten as

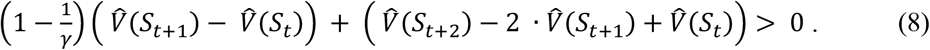

The first term on the left-hand side in Inequality (8) corresponds to the temporal derivative of estimated value, scaled by a negative constant, whereas the second term corresponds to the second derivative of estimated value (Mikhael et al.).

Inequality (8) represents the condition for ramping TD RPEs. Note that when *γ* is close to one, the first term is close to zero. Hence, the condition is approximately that the second derivative must be greater than zero, a property that all convex functions satisfy. The exact condition is more restrictive, however: Because value increases with time, the first term is negative. Hence, for the condition to be satisfied, the second term must be positive enough to outweigh the negative first term. Roughly speaking, value functions must be “convex enough” to satisfy the ramping condition.

For an illustration, in Fig. S1A we show TD RPEs corresponding to a number of different value functions. It is straightforward to show that value functions in the top panel that are drawn in green satisfy the ramping condition over the evaluated domain, whereas those drawn in red do not. As shown in the bottom panel, only value functions satisfying the ramping condition (Inequality (8)) produce ramping TD RPEs.

### Dopamine responses with speed manipulation can be captured by TD RPEs whose value functions satisfy the ramping condition

We take states to correspond to locations, and let us assume that the value function is quadratic with states (i.e., .. 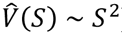). This value function obeys the ramping condition (green curves in Fig. S1A). We show in Fig. S1B how progressing through states with variable speed results in value functions of varying convexity (and first derivative) when plotted against time. Applying Equation (4) to these value functions results in TD RPEs that ramp and whose magnitudes increase with speed.

## SUPPLEMENTARY MOVIES

**Supplementary Movie 1. Visual stimulus in the standard approach-to-target condition.**

**Supplementary Movie 2. Visual stimulus in the long-distance teleport condition.**

**Supplementary Movie 3. Visual stimulus in the short-distance teleport condition.**

**Supplementary Movie 4. Visual stimulus in the pause condition.**

**Supplementary Movie 5. Visual stimulus in the fast (x2) speed condition.**

**Supplementary Movie 6. Visual stimulus in the slow (x0.5) speed condition.**

